# Genetic control of mitotic-to-meiotic transition regulates germline cell survival

**DOI:** 10.64898/2026.05.28.728457

**Authors:** Juan Garrido-Maraver, Carlos Estella, Acaimo González-Reyes

## Abstract

The coordination between DNA damage repair and cell cycle progression is essential to ensure cell survival and organ homeostasis. This is particularly critical during gametogenesis, where germline cells first proliferate and then transition from mitosis to meiosis. Meiotic cells frequently undergo recombination, which itself implies the generation of severe DNA damage in the form of double-strand DNA breaks (DSBs) that ought to be repaired to preserve genome integrity. Here, we identify *Drosophila bru1* as an essential factor in the mitotic-to-meiotic transition in the female germline. The RNA-binding protein Bru1 is a translational repressor of multiple targets, including cyclins, and is sharply upregulated at the mitotic-to-meiotic boundary. We show that loss of *bru1* disrupts germline development by altering cell cycle regulation, increasing DNA damage and cellular stress, and triggering apoptosis. *bru1* mutants display clear signs of accelerated mitotic activity leading to extra divisions in the germline, in agreement with their elevated Cyclin A and B levels. Importantly, slowing cell cycle progression in *bru1* mutants via *string/cdc25* knockdown decreases DNA damage and cell death. Mechanistically, *bru1* regulates *mei-W68* transcription, the topoisomerase responsible for DSB production in the germline. Higher Mei-W68 levels induce premature and ectopic DSBs, which persist longer in the mutant germline, indicating defective repair and potentially resulting in p53-mediated apoptosis. Our work classifies *bru1* as a safeguard of genome integrity and germline survival during the early stages of *Drosophila* female gametogenesis. *bru1* regulates Mei-W68 levels and DSB formation, and controls the mitotic-to-meiotic transition by influencing cell cycle progression. The existence of *bru1* homologues in mammals with established roles in gametogenesis suggests a broader biological relevance of our discoveries.

## Introduction

Throughout human development and growth, cells proliferate to form individuals containing 30+ trillion cells (Hatton *et al*, 2023). In adulthood, controlled cell division and differentiation maintain tissue homeostasis, with proliferation rates reaching up to 3.8 million divisions per second in a standard adult male (Sender & Milo, 2021). Importantly, cells are exposed to risks that challenge DNA quality throughout the cell cycle (reviewed in (Yousefzadeh *et al*, 2021)). In fact, it is estimated that an average of ∼10^4^ DNA-damage events occur per cell and day (De Bont & van Larebeke, 2004), of which 10-50 are DSBs (Vilenchik & Knudson, 2003). Since the stability of genomes is fundamental for organismal survival, it is essential that repair mechanisms act efficiently to preserve genome integrity, particularly so in gametogenesis, as gametes ought to ensure proper genetic transmission to offspring. Still, during gametogenesis, germ cells proliferate and then activate the meiotic programme (Mehrotra & McKim, 2006), two known sources of DNA damage in the form of replication stress-associated DNA breaks and recombination DSBs, respectively. Both types of DNA breaks require timely repair, a process that involves the activation of the p53 regulatory network (Mehrotra & McKim, 2006; Park *et al*, 2019; Chakravarti *et al*, 2022; Molano-Fernández *et al*, 2025). The tumor suppressor p53 sits in the center of the DNA damage response that safeguards genomic stability trough the activation of different cellular responses such as cell cycle arrest and DNA repair (Baonza *et al*, 2022; Vousden & Lane, 2007). When damage is irreparable, cells typically undergo p53-dependent programmed cell death (Roos & Kaina, 2006; Vousden & Lane, 2007). In *Drosophila*, p53 levels can modulate different cellular responses. High levels activate the apoptotic-regulatory genes (i.e. *hid* and *reaper*) to promote the apoptotic cascade, allowing initiator caspases such as *Dronc* trigger the executioner caspases *Drice* and *Dcp-1*, which in turn degrade cellular structures, fragment DNA, and induce cell morphology changes, ultimately leading to cell death (Fig. 1A) (Brodsky *et al*, 2000; Steller, 2008; Baonza *et al*, 2022). That germline cells are sensitive to apoptosis is demonstrated by the fact that entire germline cysts can undergo cell death from very early stages as a response to structural defects and/or environmental conditions, such as starvation (Lebo & McCall, 2021). Although p53 is activated by meiotic recombination events in *Drosophila* and vertebrates without inducing apoptosis (Lu *et al*, 2010; Wylie *et al*, 2014), the specific molecular mechanisms that allow germ cells to escape DNA damage-induced apoptosis are largely unknown.

**Figure 1.**
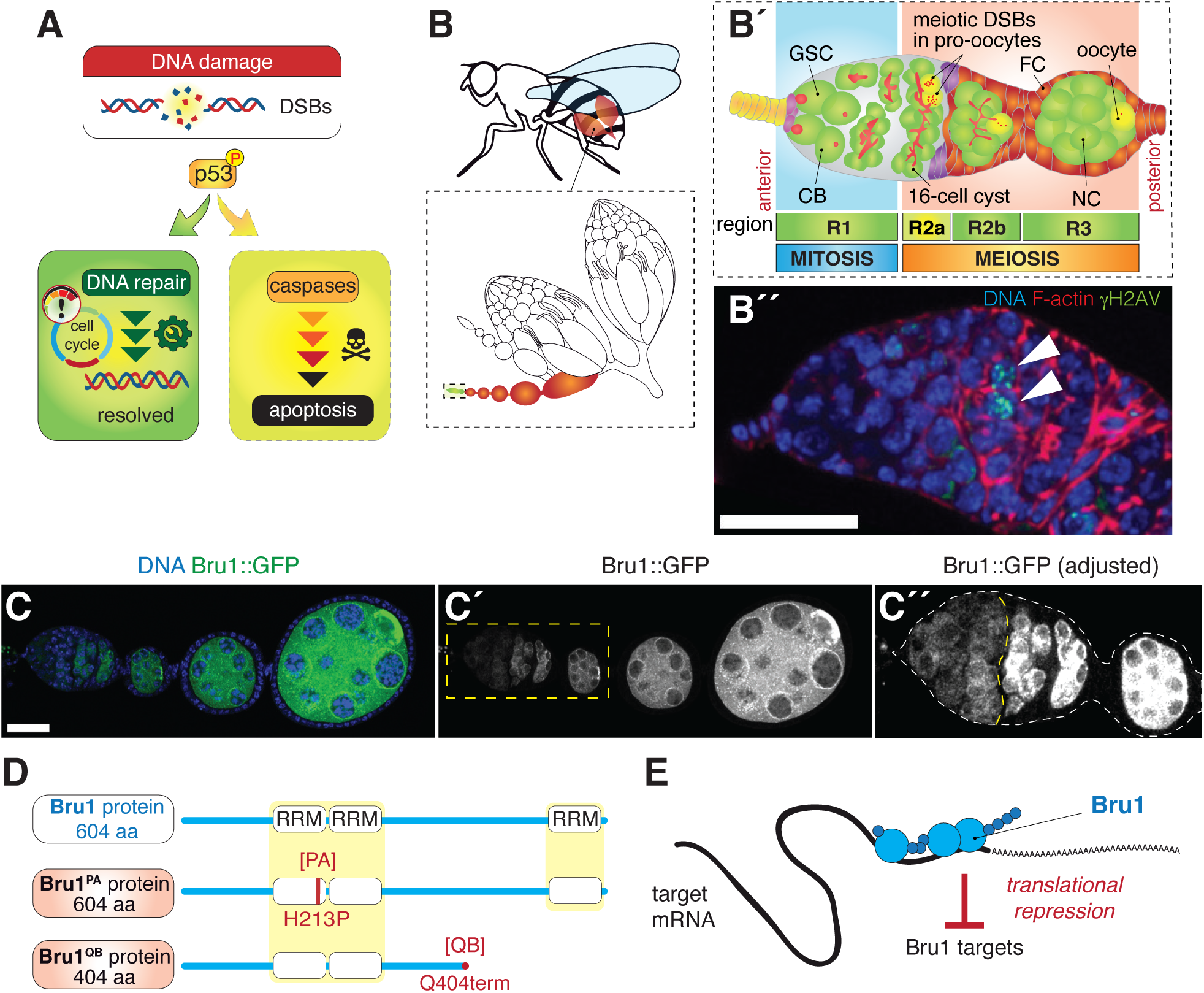
*bru1* encodes a translational repressor. (**A**) Simplified scheme of the cellular response upon DNA damage that activates DNA repair mechanisms, including cell cycle regulation. Unresolved DNA breaks can result in caspase-mediated, p53-dependent apoptosis. (**B**, **B′**) Schematic representation of the relevant cell types within a germarium. (**B′′**) Germarium from a control female stained to visualize DSBs (anti- γH2AV; green), F-actin (Rhodamine-Phalloidin; red) and DNA (Hoechst; blue). White arrowheads point to the pro-oocytes. (**C, C’, C′′**) Ovariole showing the endogenous pattern of expression of *bru1* (Bru1::GFP, green) and DNA (Hoechst, blue). Yellow dashed line in C′′ marks the mitotic-to-meiotic transition boundary. Levels of Bru1::GFP signal were adjusted in C′′ to see region 1 in detail. (**D**) Scheme of Bru1 protein, isoform A, highlighting the RNA recognition motives (RRM). The molecular nature of the PA and QB mutations is indicated. (**E**) Scheme of Bru1 as a translational repressor. GSC, germline stem cell; CB, cystoblast; FC, follicle cell; NC, nurse cell; DSBs, double-strand breaks; aa, amino acid. Scale bars: 25 μm.

During *Drosophila* oogenesis, gamete production starts in the germarium, a tapered structure found at the anterior tip of each ovariole, the egg producing tubules that conform the ovaries. It is in the germarium where germ cells first proliferate and later enter meiosis. Mitoses take place in region 1 of the germarium, where germline stem cells (GSCs) divide asymmetrically giving rise to daughter GSCs and to differentiating cystoblasts (CBs; Fig. 1B). CBs undergo four rounds of incomplete divisions to produce 16-cell cysts that enter region 2. In this region, further subdivided into regions 2a and 2b, the entire cyst is encapsulated by follicle cells (Lin & Spradling, 1993). Importantly for this work, germline cells transition from mitosis to meiosis in region 2. Key steps in this transition include the downregulation of the mitotic Cyclins (A and B) (Sugimura & Lilly, 2006; Bayer *et al*, 2025) and the activation of the meiotic programme, evidenced by the appearance of recombination nodules and the formation of the synaptonemal complex in several cystocytes per cyst, including the two pro-oocytes (the first cystocytes to form, from which the future oocyte will be selected; reviewed in (Hughes *et al*, 2018; Lake & Hawley, 2012)). Meiosis in the *Drosophila* female implies the formation of a discrete number of recombination-induced DSBs through the activity of the Mei-W68 topoisomerase and additional factors such as chromosome-associated protein Mei-P22 (Liu *et al*, 2002; Hughes *et al*, 2018; McKim & Hayashi-Hagihara, 1998). These DSBs, which can be visualized with the γH2AV marker, are largely repaired in regions 2 and 3 (Lake *et al*, 2013). Region 3 is characterized by oocyte specification and by the change in shape of the 16-cell cysts, which switch from a lens-shape arrangement to a sphere with the newly specified oocyte now placed at the posterior (Fig. 1B) (De Cuevas & Spradling, 1998; Godt & Tepass, 1998; González-Reyes & St Johnston, 1994).

Considering its molecular activity, its mRNA targets and its mutant phenotypes, we focused on the functions of *bruno 1* during the transition from mitosis to meiosis in female gametogenesis. *bruno 1 (bru1)*, also known as *bruno, arrest* or *aret*, has an essential function during oogenesis as mutant alleles disrupt oogenesis at different stages, leading to egg chamber degeneration and eventually to egg failure (Parisi *et al*, 2001; Schupbach & Wieschaus, 1991; Sugimura & Lilly, 2006; Webster *et al*, 1997). *bru1* belongs to an evolutionarily conserved gene family that is highly expressed in the *Drosophila* germline (Fig. 1C) and in somatic tissues (Good *et al*, 2000). It encodes a series of transcripts and isoforms that share structural features, particularly RNA recognition motifs, which are key to its function (Fig. 1D; Fig. S1A, B). Bru1 acts as a translational repressor that binds Bruno Response Elements (BREs; specific sequences often found in the 3’UTR of messenger RNAs; Fig. 1E), rendering them susceptible to translational repression (Kim-Ha *et al*, 1995; Webster *et al*, 1997). Among others, including *oskar, gurken* and *Sex-lethal*, a typical target of Bruno is mitotic *cyclin A*. Cyclin B levels are also increased in *bru1* mutant germ cells but it is unknown if Bru1 targets directly *cyclin B* mRNA (Wang & Lin, 2007; Sugimura & Lilly, 2006; Filardo & Ephrussi, 2003; Chekulaeva *et al*, 2006; Kim-Ha *et al*, 1995; Bayer *et al*, 2025). We demonstrate that, early in oogenesis, *bru1* contributes to the mitosis-to-meiosis transition and prevents excess DNA damage and apoptosis in the germline. Our results thus reveal an essential function for *bru1* in the protection of germline integrity through the regulation of Mei-W68 topoisomerase levels.

## Results

### Loss of *bru1* function causes mitotic cell cycle defects in germarial stages

In *Drosophila*, the mitotic cyclins Cyclin A and Cyclin B and the String phosphatase activate the G2/M promoting factor Cdk1 and regulate cell cycle progression and entry into mitosis (Lehner & O’Farrell, 1990). During early oogenesis, Cyclin A and Cyclin B are expressed in mitotic region 1 and disappear shortly after meiosis entry in early region 2 (Lilly *et al*, 2000). It is known that the translational repressive function of *bru1* targets *Cyclin A* and that both Cyclin A and B are up-regulated in *bru1* mutant germline (Sugimura & Lilly, 2006; Bayer *et al*, 2025). In fact, *bru1* is upregulated at the mitotic/meiotic boundary in region 2 of the germarium, precisely where cyclin levels decrease (Fig. 1C). Considering the role of mitotic Cyclins in cell cycle progression, we wished to determine if *bru1* mutants displayed abnormal mitotic cell cycle regulation in the germline in region 1. We utilized a viable combination of weak (*bru1^PA^*) and strong (*bru1^QB^*) alleles to generate a loss-of-function situation. First, we quantified the number of cells in DNA replication (S) phase using EdU as a marker for DNA synthesis in the germarium. Compared to controls, we observed a significantly higher number of S-phase cells in *bru1^PA/QB^* mutants in germarial region 1 (control: 1.51±0.22, n=41 germaria analyzed; *bru1^PA/QB^*: 3.37±0.42, n=35; Fig. 2A-C). Second, we scored cells in mitosis using the phospho-histone 3 (PH3) marker and found again a significant increase in mitotic cells in region 1 of *bru1^PA/QB^*mutants compared to controls (control: 0.36±0.12, n=28 germaria analyzed; *bru1^PA/QB^*: 1.07±0.20, n=28; Fig. 2D-F). The higher number of cells undergoing active mitosis might be explained by an acceleration of the cell cycle in mutant cysts, which would then divide more frequently. In such scenario, we should detect an increase in the number of cysts in mutant germaria. In fact, we observed a 40% increase in the number of cystoblasts and cysts in region 1 of *bru1^PA/QB^* mutants compared to controls (control: 5.19±0.57, n=21 germaria analyzed; *bru1^PA/QB^*: 7.28±0.46, n=18; Fig. 2G). Importantly, this significant rise in mutant cyst numbers (which is most likely an underestimation, as a considerable number of mutant cysts are lost to apoptosis and are not thus included in the data set; see next section) could not be explained by a similar increase in GSCs, as we detected only a 10% increase in the latter (control: 2.40±0.08, n=70 germaria analyzed; *bru1^PA/QB^*: 2.64±0.09, n=53; Fig. 2H). This finding further supports the idea of mitotic cell cycle acceleration in *bru1^PA/QB^* mutant germaria. Third, we made use of the Flyfucci tool (Zielke *et al*, 2014), which allows classification of cells into the G1, S and G2/M phases. We quantified RFP and GFP nuclear signals from cells in germarial region 1 and calculated RFP/GFP ratios (Fig. 2I). Based on reporter signal intensities and threshold criteria (see Methods for details), we classified cells according to their cell cycle phase (Fig. 2J). We found a 32% increase in the proportion of S-phase *bru1^PA/QB^* mutant cells, in agreement with our EdU data, and a 43% rise in the fraction of G1 mutant cells (n=10). Concomitantly, the percentage of cells in G2/M was decreased. The increased percentages of mutant germ cells in G1 or S are consistent with both the prolonged activation of Cdk1 (which we interpret as a consequence of elevated Cyclin A and B levels) and the acceleration of the G2 phase of the cell cycle.

**Figure 2.**
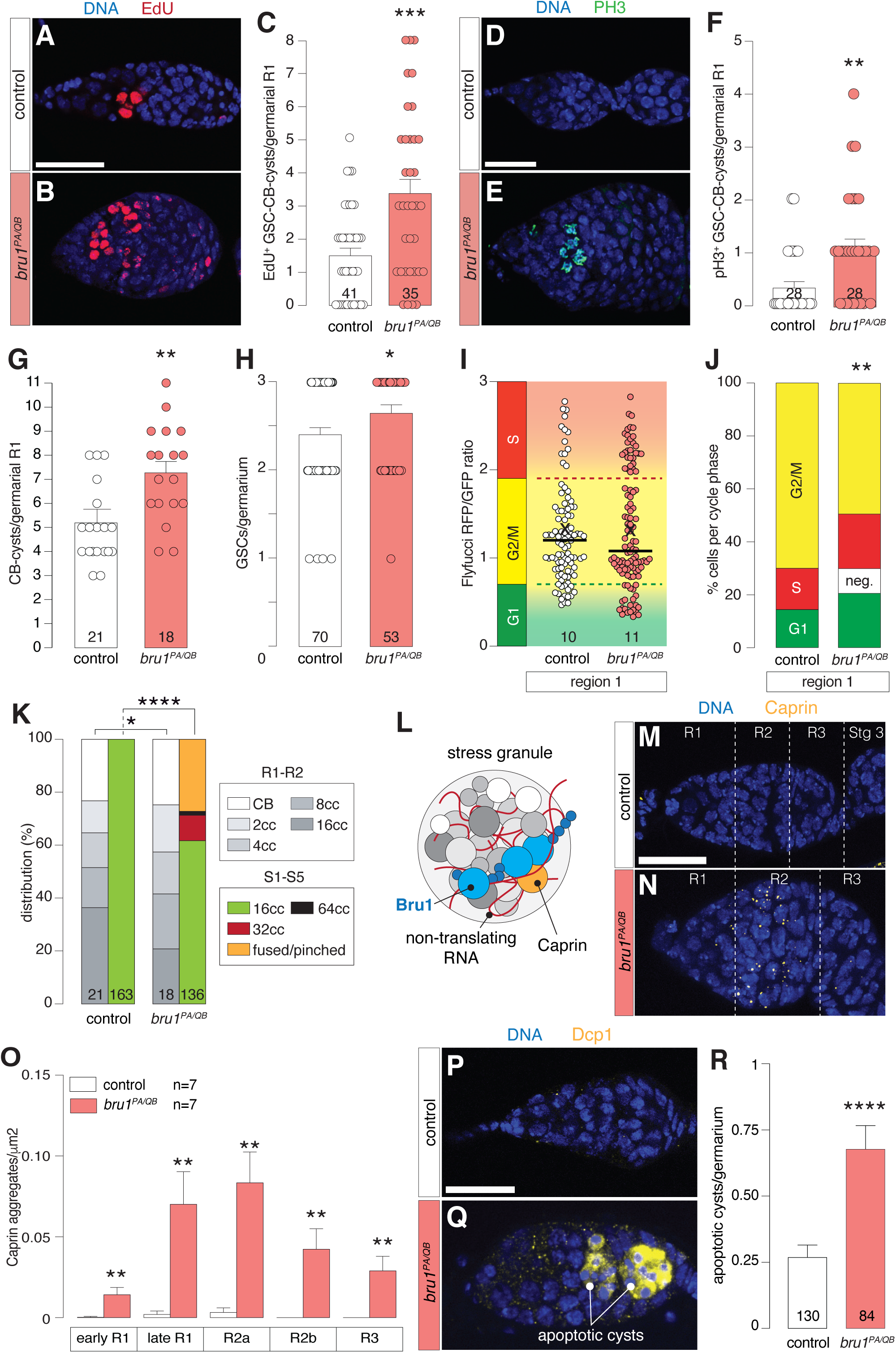
Loss of *bru1* function causes mitotic cell cycle defects, stress granule formation and apoptosis activation in germarial stages. (**A, B**) Germaria stained to visualize S-phase cells (EdU, red) and DNA (Hoechst, blue). (**C**) Quantification of the number of EdU^+^ GSC, CB and 2-8 cell cysts per germarial region 1. (**D, E**) Germaria stained to visualize mitotic cells (anti-PH3, green) and DNA (Hoechst, blue). (F) Quantification of the number of PH3^+^ GSC, CB and 2-8 cell cysts per germarial region 1. (**G**) Quantification of the total number of CB and 2-8 cell cysts per germarial region 1. (**H**) Quantification of the total number of GSCs per germarium. (**I**) Flyfucci signals obtained per cell in region 1 shown as the RFP/GFP ratio. Red and green dashed lines represent thresholds to define cell cycle phases (G1, G2/M and S; see Methods for details). (**J**) Percentages of cells classified into different cell cycle phases. (**K**) Distribution (%) of CB and 2-16 cell cyst, of egg chambers (S1-5) containing 16, 32 or 64 germ cells and of fused/pinched egg chambers. (**L**) Scheme of a stress granule containing non-translating RNAs and multiple proteins, including Caprin and Bru1. (**M**-**N**) Germaria stained to visualize stress granules (anti-Caprin, yellow) and DNA (Hoechst, blue). (**O**) Quantification of the number of stress granules per area though germarial regions. (**P**-**Q**) Germaria stained to visualize apoptosis activation (anti-Dcp1, yellow) and DNA (Hoechst, blue). (**R**) Quantification of the number of apoptotic cysts per germarium. Statistical significance was calculated with unpaired Student’s (C, F, G, H, O, R), Mann-Whitney unpaired nonparametric (I), and Chi-square t-tests (J, K). Scale bars: 25 μm.

The above results indicate that the *bru1^PA/QB^* mutants exhibit cell cycle defects in the germarium. Quantification of the germline in controls and *bru1^PA/QB^* mutants revealed that the latter, while maintaining a similar distribution of cystoblasts and 2- to 16-cell cysts, produced a considerable number of 32- and ∼64-cell cysts, consistent with additional rounds of mitosis. Furthermore, we detected a substantial percentage of cysts with abnormal germ cell numbers arising either from egg chamber fusions (resulting in germ cell counts that are multiple of 16) or from defective egg chamber budding in germarial region 3 (where the combined contents of adjacent cysts, each with fewer than 16 cells, totaled 16; Fig. 2K; Fig. S2A, B). These phenotypes likely reflect a lack of coordination between germline cyst development and the proliferation and migration of somatic follicle cells responsible for encapsulating the germline in region 3. Such discoordination could result from cell cycle irregularities in *bru1* germline mitoses. In summary, we conclude that the mutant germline possesses an accelerated cell cycle and divides more frequently, as indicated by the increase in germline cells, EdU and PH3 labeling, by the appearance of cysts with extra mitoses and by the aberrant expression of *cyclin A* and *B* in *bru1* mutant germaria (Bayer *et al*, 2025; Sugimura & Lilly, 2006).

### Loss of Bru1 function induces stress response and apoptosis in germarial stages

*bru1*-dependent degeneration was originally associated with a high frequency of necrosis, indicating the requirement of *bru1* for cell survival (Parisi *et al*, 2001). To determine whether the absence of *bru1* induced a stress response in the germarium, we analyzed the distribution of the stress granule component Caprin (Solomon *et al*, 2007) in control and *bru1^PA/QB^* trans-heterozygotes. We found that the mutant combination displayed a significant increase in the number of stress granules in germarial regions 1 and 2, thus confirming that the absence of *bru1* function induces a stress response in the germline (n=7 germaria analyzed; Fig. 2L-O). Next, we explored cell death events and found that *bru1* mutants exhibited increased apoptosis, as measured by the activation of the effector caspase Dcp1. The most degenerated egg chambers in both hemizygous and trans-heterozygous mutants displayed high levels of active (cleaved) Dcp1 and pyknotic nuclei affecting both germline and somatic tissues (Fig. S3C-G). Notably, we also detected apoptosis activation within the germarium, with mutant tissues consistently showing higher frequencies of apoptotic cysts in region 2 compared to controls (control: 0.27±0.05, n=130 cysts analyzed; *bru1^PA/QB^*: 0.68±0.09, n=84; Fig. 2P-R; Fig. S3G).

### *bru1* activity is required for proper DSB repair

Given the increased mitotic activity in mutant germline, the augmented levels of cellular stress and death observed in *bru1* mutant germaria, and considering the relationship between apoptosis, formation of stress granules and DNA lesions (Byrd *et al*, 2016; Verkaik *et al*, 2008; Singatulina *et al*, 2022), we investigated whether *bru1* mutants displayed abnormal levels of DNA damage. In the germarium, meiotic recombination of homologous chromosomes courses with the controlled induction and repair of DNA breaks (McKim & Hayashi-Hagihara, 1998; Jang *et al*, 2003). To assess whether *bru1* mutant germaria exhibited abnormal DSB production, we used an antibody that recognizes phosphorylated H2AV (anti-γH2AV). H2AV phosphorylation is considered an early marker of checkpoint activation, mediated by ATM and ATR, upon DNA damage, replication stress or meiotic recombination (Molano-Fernández *et al*, 2025; Joyce & McKim, 2009). In meiotic cells, γH2AV is commonly used to mark programmed DSBs (Lake *et al*, 2013). We analyzed nuclear γH2AV levels in germline cells within the germarium. As expected and consistent with the programmed DSBs and their subsequent repair in meiotic cells (Lake *et al*, 2013; Mehrotra & McKim, 2006), control germaria exhibited a peak of γH2AV signal in region 2a, which decreased in regions 2b and 3 (Fig. 3A). Next, we calculated γH2AV-positive signal intensity in different *bru1* mutant backgrounds and observed higher values in most stages analyzed including region 3 (Fig. 3B-C; Fig. S3H, I). In fact, when analyzed in detail, the intensity of the nuclear signal in region 2a and 2b of control and *bru1^PA/QB^* germaria indicated remarkable differences: 30% of the values in region 2a and 75% in region 2b of *bru1^PA/QB^* germaria were above the 99^th^ percentile values of the equivalent control levels (Fig. 3D). We then scored the number of γH2AV foci in region 2 pro-oocytes of control germaria, obtaining values in the same range as previously reported (Mehrotra & McKim, 2006; Lake *et al*, 2013). In contrast, *bru1^PA/QB^* mutant pro-oocytes showed higher numbers of γH2AV foci compared to controls (mean number of γH2AV foci in region 2a: control=11.43±0.61, n=28 cystocytes analyzed; *bru1^PA/QB^*=13.19±0.73, n=26; Fig. 3E-G). Overall, these data indicate that *bru1* mutants exhibit an abnormal accumulation of DSBs during oogenesis that are not resolved in region 3, which may be linked to the increased apoptosis observed in mutant germaria. Interestingly, while in control region 2b cysts the number of cells containing high levels of nuclear γH2AV signal was most often two (likely the pro-oocytes, n=59 cysts analyzed), *bru1^PA/QB^* germaria showed increased numbers of cystocytes with high γH2AV amounts in region 2 cysts (4.47±0.30 cells per cyst, n=78; Fig. 3H). Thus, considering that mutant germ cells contain more γH2AV foci and higher levels of γH2AV signal, and that other cells in addition to the two pro-oocytes display visible DNA damage, we surmise that *bru1* activity normally prevents excess, unspecific DSB formation. Finally, we scored the presence of DSBs in region 1 mitotic germline cells. As expected, we hardly detected phosphorylated H2AV foci in controls, while *bru1* mutants displayed significantly higher numbers of γH2AV-positive events (Fig. 3I-K). While this result may also be interpreted as a precocious entry into meiosis of *bru1* region 1 cysts —hence the higher levels of Mei-W68 protein and the appearance of region 1 DSBs—, the similar pattern of expression of the synaptonemal complex component C(3)G in control and mutant germaria indicated that meiosis was properly timed in mutant germ cells (Fig. S4A-C).

**Figure 3.**
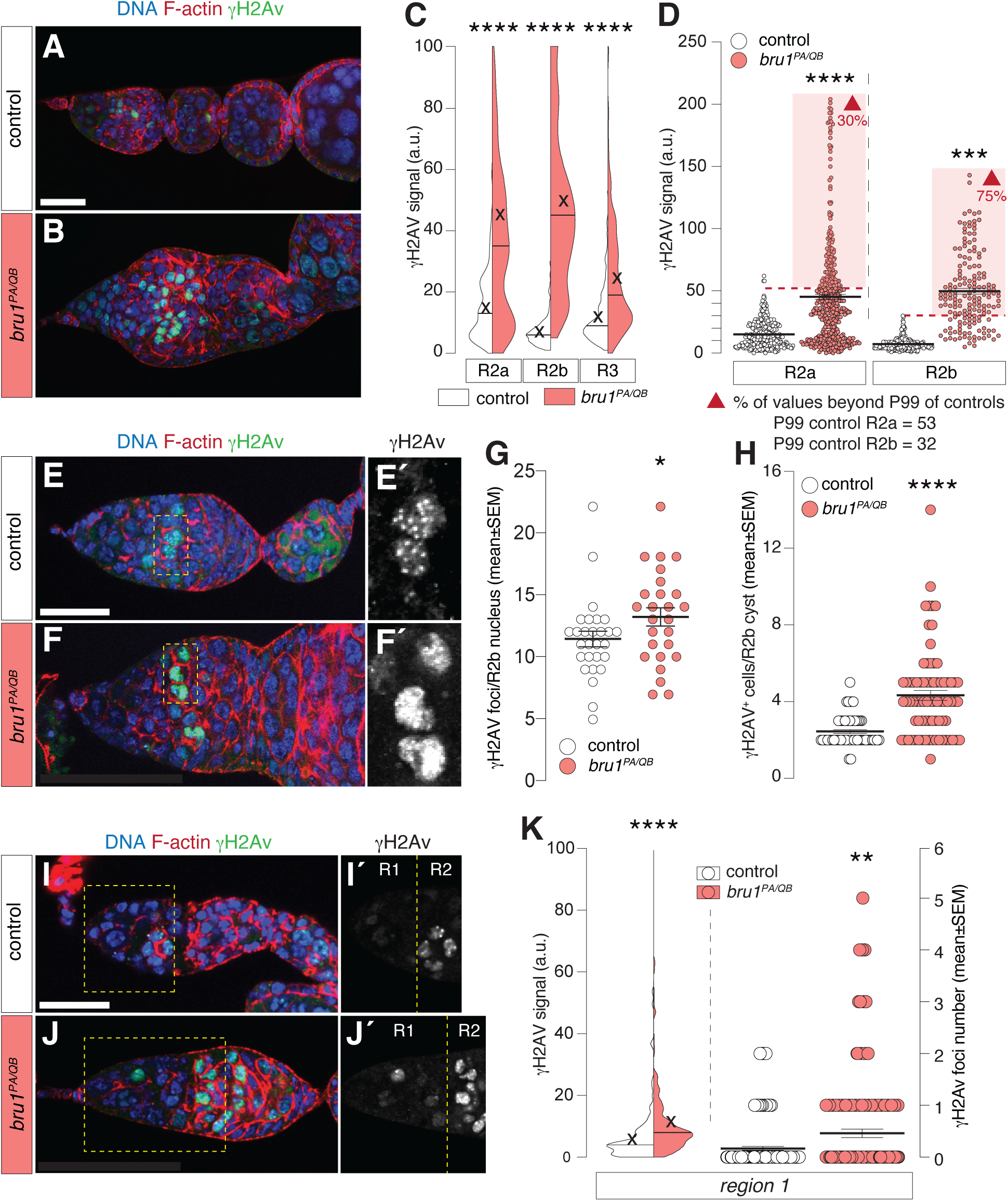
Proper DSB repair in the germline requires *bru1* activity. (**A, B**) Germaria stained to visualize DSBs (anti-γH2AV, red) together with F-actin (Rhodamine: Phalloidin, red) and DNA (Hoechst, blue). (**C**) Quantification of nuclear γH2AV signal in regions 2a, 2b and 3. (**D**) Scatter plot representing individual γH2AV signal values in regions 2a and 2b. Red, dashed lines mark the 99^th^-percentile threshold of controls. In the mutant samples, 30% of values in region 2a and 75% in region 2b exceed this control threshold. (**E**, **F**) Germaria stained to examine the F-actin (Rhodamine: Phalloidin, red), γH2AV (anti-γH2AV, green) and DNA (Hoechst, blue) signals in region 2. Insets show γH2AV foci and signal intensity in region 2b. (**G**) Quantification of the number of γH2AV foci in region 2b pro-oocytes. (**H**) Quantification of the number of γH2AV^+^ cells per region 2b cyst. (**I**, **J**) Detail of region 1 in germaria stained to visualize DSBs (anti-γH2AV, red). (**K**) Quantification of the number of γH2AV nuclear signal and foci number in region 1 germline cells. Statistical significance was calculated with a Mann-Whitney unpaired nonparametric t-test (C, D, G, H, K). Scale bars: 25 μm.

Next, we wished to ascertain whether it was DSB accumulation what led to the increased apoptosis characteristic of *bru1* mutant germaria. We generated recombinant chromosomes carrying both *bru1* and *mei-W68* mutant alleles. *mei-W68* encodes the topoisomerase II-like involved in the generation of DSBs in region 2. Hence, *mei-W68* mutant females show a remarkable decrease in meiotic DSBs (McKim & Hayashi-Hagihara, 1998). Double mutant females *bru1 mei-W68* displayed a strong reduction in γH2AV levels in the germarium –including region 1–, and completely abolished the induction of apoptotic cysts in the germarium when compared to the single *bru1* mutant (Fig. 4A-E). In addition, we compared the percentage of cystoblasts and region 1 cysts positive for EdU in *bru1* mutants and *bru1 mei-W68* double mutants, and found no significant differences, suggesting that the increase in DNA damage in *bru1* mutants might be primarily linked to the catalytic function of Mei-W68 and not to the enhanced mitotic activity of the mutant germ cells (Fig. 4F; Fig. S4D, E).

**Figure 4.**
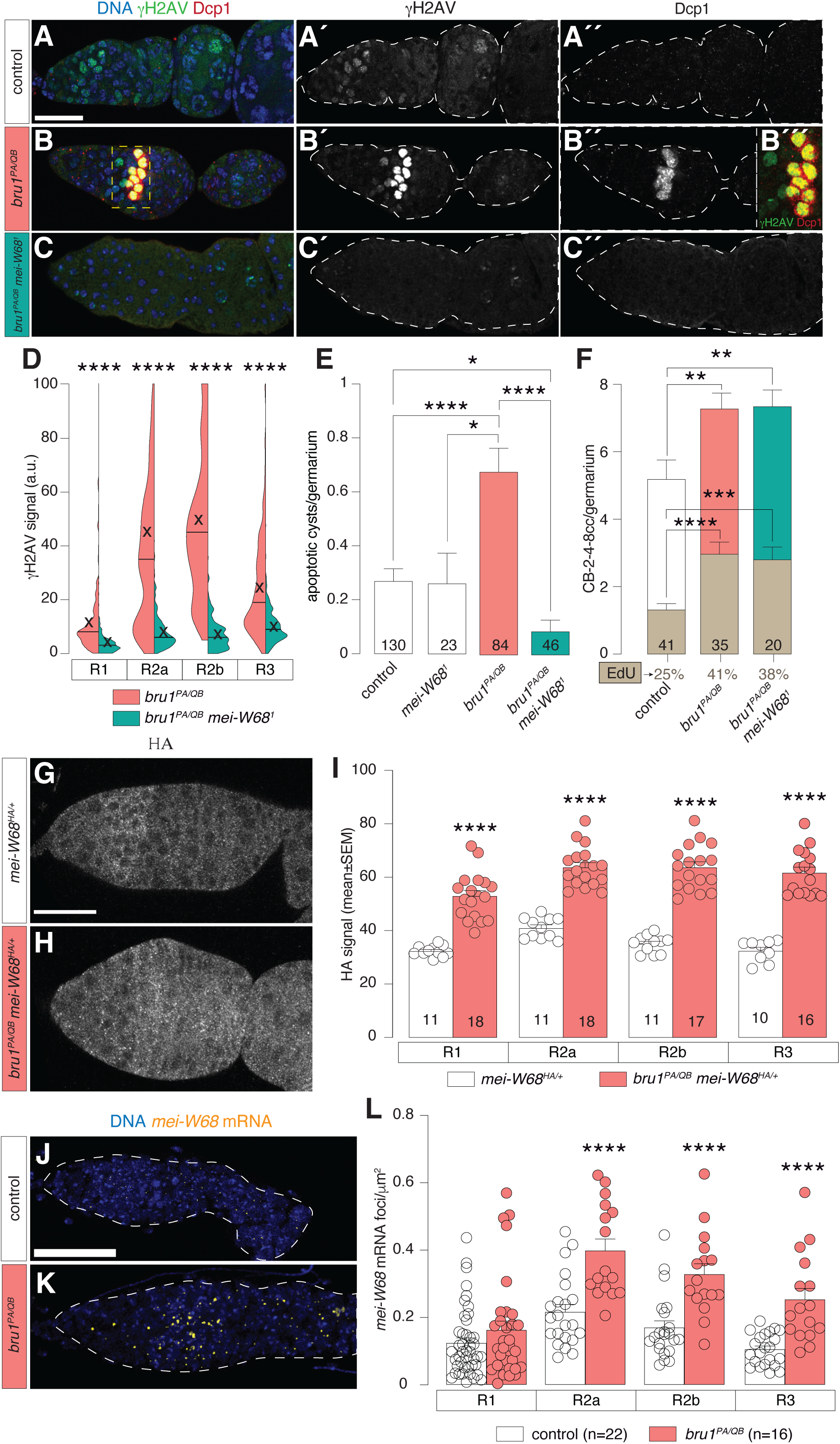
DSB accumulation drives apoptosis in *bru1* mutant germaria. (**A**-**C**) Germaria stained to visualize DSBs (anti-γH2AV, green), apoptosis (anti-Dcp1, red) and DNA (Hoechst, blue). Inset in (B′′′) shows colocalization of γH2AV and Dcp1 signals. (**D**) Quantification of nuclear γH2AV signal in regions 1, 2a, 2b and 3. (**E**) Quantification of the number of apoptotic cysts per germarium. (**F**) Quantification of the total number of GSCs, CBs and 2-8 cell cysts in controls and experimental conditions. Bars in gold indicate number of EdU^+^ GSCs, CBs and cysts. The percentage of EdU^+^ *versus* total is shown below for each genotype. (**G**, **H**) Germaria stained to visualize HA-tagged Mei-W68 (HA, green) together with F-actin (Rhodamine: Phalloidin, red) and DNA (Hoechst, blue). (**I**) Quantification of HA signal in regions 2, 2a, 2b and 3. (**J**, **K**) Germaria stained to visualize *mei-W68* mRNA (probe, yellow) together with DNA (Hoechst, blue). (**L**) Quantification of HCR-positive foci per area in regions 1, 2a, 2b and 3. Statistical significance was calculated with Mann-Whitney unpaired nonparametric (D) and unpaired Student’s t-tests (E, F, I, L). Scale bars: 25 μm.

To investigate the origin of ectopic region 1 DSBs and of excess DSBs in region 2 in mutant germaria, we quantified endogenous HA-tagged Mei-W68 (Vallés *et al*, 2024) levels and found that *bru1* mutants showed significantly increased levels compared to controls in all germarial stages (Fig. 4G-I; Fig. S4F). In addition, direct visualization of *bru1* mRNA utilizing hybridization chain reaction (HCR) confirmed that *bru1* controlled *mei-W68* transcription. First, mutant germaria exhibited a significantly higher number of HCR-positive foci than controls (Fig. 4J-L), and the average size of the HCR-positive dots was also significantly increased in mutant conditions (Fig. S4G). Since HCR yields semi-quantitative measurements of transcript abundance, these results strongly suggest that *mei-W68* transcription is upregulated in *bru1* mutant cells. In all, our findings reveal that the abnormal accumulation of DSBs in *bru1* mutants i) depends on increased *mei-W68* transcription and elevated Mei-W68 levels and ii) directly underlies the excess apoptosis observed in experimental germaria. Furthermore, the fact that the mitotic phenotypes and the excess DSBs arise before stress granules indicates that the latter is likely a consequence of the former.

### *bru1* acts in the germline for cyst survival and DSB repair

To determine if *bru1* was required in the germline, we down-regulated *bru1* specifically in the germline using the *nanos-Gal4* (*nos-Gal4*) driver and an RNA interference line (*bru1 RNAi*). *nos>bru1 RNAi* ovarioles exhibited a near-complete loss of ovarian structural integrity and were sterile. Importantly, they showed increased cellular stress and DNA damage in the germline as indicated by Caprin and γH2AV stainings and by the presence of apoptotic cysts, thus closely mimicking the *bru1* mutant phenotypes (Fig. S5A-H). Conversely, overexpression of the same RNAi line with the somatic cell driver *traffic jam-Gal4 (tj-Gal4)*, did not give *bru1* phenotypes in adult germaria (Fig. S5I-K). We conclude that, at least in the germarium, *bru1* acts fundamentally in the germline to control cyst survival and proper DNA damage production/repair.

We have found that *bru1* mutant region 1 germ cells displayed DSBs (Fig. 3I-K). This unexpected result indicated that *bru1* prevented untimely, *mei-W68*-dependent DNA damage in mitotic cysts. To confirm this, we made use of the *bam-Gal4* line, expressed in region 1 CBs and early cysts (McKearin & Spradling, 1990), to remove *bru1* function in mitotic region 1. Quantification of the number of γH2AV foci per region 1 cell of control and *bam>bru1 RNAi* samples detected a significant increase of foci in the mutant condition (control: 0.17±0.09 (n=79); *bam>bru1 RNAi*: 0.97±0.15 (n=102); Fig. S5L-O). This is consistent with a recent transcriptomic analysis of germline differentiation, which described the expression of *bru1* in region 1 (Samuels *et al*, 2024) and with the pattern of expression of Bru1::GFP.

### Sustained mitoses in *bru1* mutant germline cells induce excess DNA damage and apoptosis

The presence of excess γH2AV-positive lesions in proliferating germ cells opens the possibility that a proportion of the DNA damage observed in mutant germaria originated independently of meiosis. We thus set out to determine whether the increase in DNA damage and apoptosis in *bru1* mutant germaria could result, at least partially, from sustained mitoses. To this end, we expressed a *string RNAi* construct in the germline and examined its impact on DSBs and apoptosis. Most cell types enter mitosis through a progressive activation of Cyclin-Cdk1 complexes. In *Drosophila,* the String phosphatase promotes Cdk1 dephosphorylation to trigger mitotic entry (O’Farrell, 2001; Di Talia & Wieschaus, 2012). Thus, removing partly *string* function delays entry into mitosis, increasing cell cycle length and allowing longer times for DNA-damage surveillance and repair. Compared to *nos>bru1 RNAi* controls, *nos>bru1 RNAi* + *string RNAi* showed a partial, albeit significant, reduction in the number of DSBs and of apoptotic cysts in the germarium (Fig. 5A-D). Moreover, knocking down *string* levels in the germline rescued significantly the number of 16-cell cysts and it largely corrected the irregularities (fusions and aberrant budding) observed during mutant egg chamber individualization (*nos>bru1 RNAi* + *string RNAi*; Fig. 5E). Altogether, our findings support a model in which loss of *bru1* function increases Mei-W68 levels in germ cells, generating non-meiotic DSBs in mitotically active region 1 cells and excess meiotic-DSBs in region 2. In addition, the sustained mitotic cell cycles in mutant germaria generate cellular stress(es), overwhelming the repair machinery and resulting in DNA damage-mediated apoptosis of mutant cysts.

**Figure 5.**
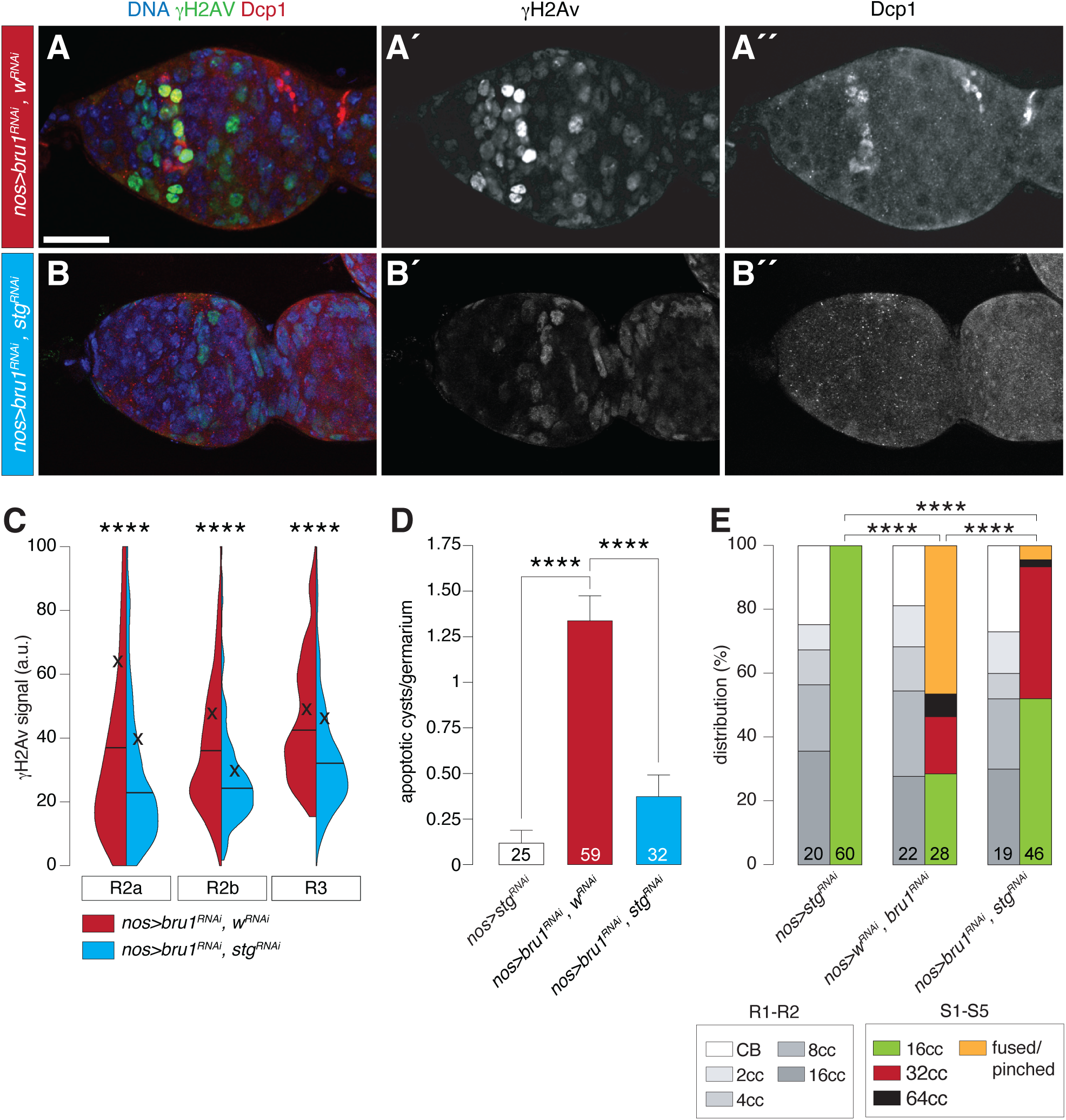
Sustained mitoses in *bru1* mutant germline induce excess DNA damage and apoptosis. (**A, B**) Germaria stained to visualize DSBs (anti-γH2AV, green), apoptotic cells (anti-Dcp1, red) and DNA (Hoechst, blue). (**C**) Quantification of nuclear γH2AV signal in regions 2a, 2b and 3. (**D**) Quantification of the number of apoptotic cysts per germarium. (**E**) Distribution (%) of CB and 2-16 cell cyst, and egg chambers (S1-5) containing 16, 32 or 64 germ cells and of fused/pinched egg chambers. Statistical significance was calculated with unpaired Student’s (C), Mann-Whitney unpaired nonparametric (D), and Chi-square t-tests (E). Scale bars: 25 μm.

### *bru1* activity in the germline prevents p53-induced apoptosis

As introduced above, unrepaired DNA damage triggers p53-dependent programmed cell death. In *Drosophila*, p53 recognizes responsive elements in apoptotic-regulatory genes such as *hid* and *reaper* to promote apoptosis (Ruiz-Losada *et al*, 2022; Vousden & Lane, 2007; Baonza *et al*, 2022). In addition, *corp*, a negative regulator of p53 in *Drosophila*, suppresses DNA damage-induced apoptosis (Chakraborty *et al*, 2015). To confirm that the increase in apoptotic cysts in the germarium depended on p53 and the subsequent induction of the apoptotic cascade, we first wished to test apoptosis in the absence of *bru1* and *p53*. However, *bru1 p53* double mutant adult females could not be obtained. We then overexpressed *corp* or downregulated *p53*, *hid* or the three pro-apoptotic genes *hid, reaper* and *grim* (utilizing the *miRHG* tool (Siegrist *et al*, 2010)) in a *bru1*-silenced background. In all instances, *nos>bru1 RNAi* + *p53 RNAi*, *nos>bru1 RNAi* + *hid RNAi, nos>bru1 RNAi* + *miRHG* and *nos>bru1 RNAi* + *corp* resulted in a significant reduction in the number of apoptotic cysts compared to control *nos>bru1 RNAi* + *w RNAi* (Fig. 6A-E). These results suggest that the apoptosis observed in *bru1* mutant germaria is, at least partially, mediated by p53 and apoptotic regulatory elements. To confirm this, we analyzed directly *hid* and *reaper* expression in the germarium using HCR and observed a notable increase in the transcription of both genes in the germline of *bru1* mutant germaria compared to controls (Fig. 6F-H). From these results we conclude that, upon *bru1* loss of function, mutant ovaries activate *p53, hid* and *rpr* to initiate the canonical apoptotic pathway.

**Figure 6.**
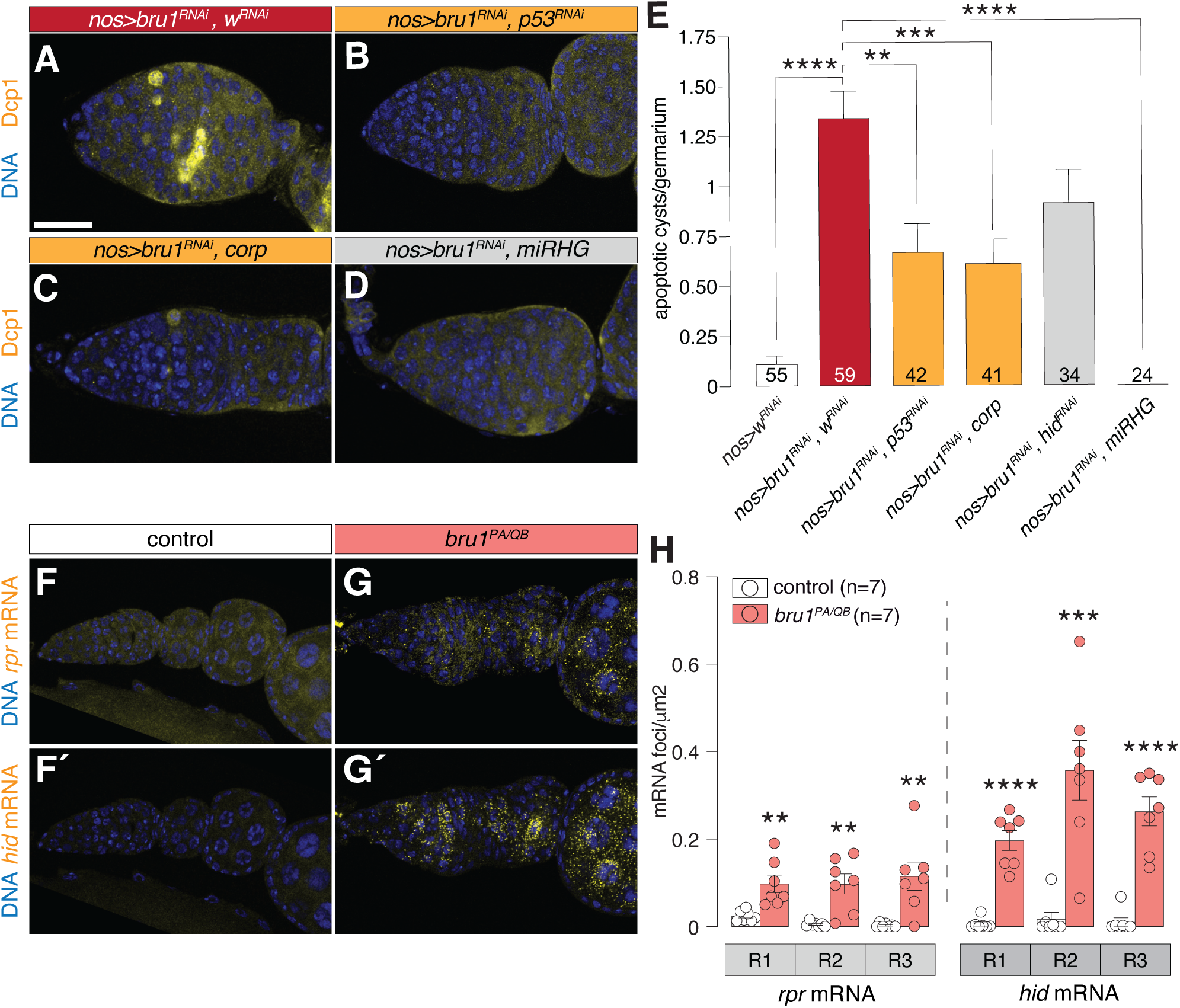
*bru1* activity in the germline prevents p53-induced apoptosis. (**A**-**D**) Germaria stained to visualize apoptosis activation (anti-Dcp1, yellow) and DNA (Hoechst, blue). (**E**) Quantification of the number of apoptotic cysts per germarium. (**F, G**) Germaria hybridized to detect mRNA of the apoptotic genes *reaper* and *hid*, and counterstained to visualize DNA (Hoechst, blue). (**H**) Quantification of the number of *hid^+^* and *rpr^+^* mRNA foci per μm^2^. Statistical significance was calculated with an unpaired Student’s t-test (E, H). Scale bars: 25 μm.

## Discussion

### *bru1* contributes to the mitosis-to-meiosis transition

Our results confirm that *bru1* is a regulator of the pace of cyst generation in the proliferating region of the germarium. During oogenesis, cysts that reach region 2 downregulate Cyclin A and B amounts, exit the mitotic mode and activate the meiotic program characteristic of gametogenesis. In contrast, *bru1* mutant cysts fail to terminate mitosis correctly and may contain 32 or 64 germ cells, an indication of supernumerary mitoses. In our model (Fig. 7), *bru1* acts as a translational repressor downregulating Cyclin A and B levels in region 2 (where Bru1 levels are highest), thus allowing the proper transition from mitosis to meiosis in female gametogenesis. The sequential progression from mitosis to meiosis in *Drosophila* involves a series of genes related to germline proliferation and differentiation, including *nos, bam*, *mei-P26* and *Sxl* (Chen *et al*, 2014; Insco *et al*, 2012; Li *et al*, 2013; Page *et al*, 2000; Benner *et al*, 2024). Importantly, beyond cyclins, *bru1* also targets *Sxl* (Wang & Lin, 2007), a splicing factor required for germline differentiation that is expressed in GSCs, CBs and early cysts (Chau *et al*, 2009). Sxl regulates splicing of multiple mRNAs including *nos*, *bam* or *mei-P26* (revised in (Mercer *et al*, 2021)), raising the possibility that *bru1* influences the shift from mitosis to meiosis not only through its effects on Cyclin A and B levels but also by modulating Sxl amounts. Because transcription in meiosis can be strongly silenced, it is therefore not surprising that RNA-binding proteins regulate the initial steps into meiosis by controlling post-transcriptionally the production of meiotic proteins. This is especially critical in the *Drosophila* female germline, in which transcriptomic analyses have revealed that the expression of genes required for female meiosis I can be regulated at the level of translation (Vallés *et al*, 2024). It would be interesting to determine whether the repressive function of Bru1 participates in the translational regulation of early-expressed meiotic genes.

**Figure 7.**
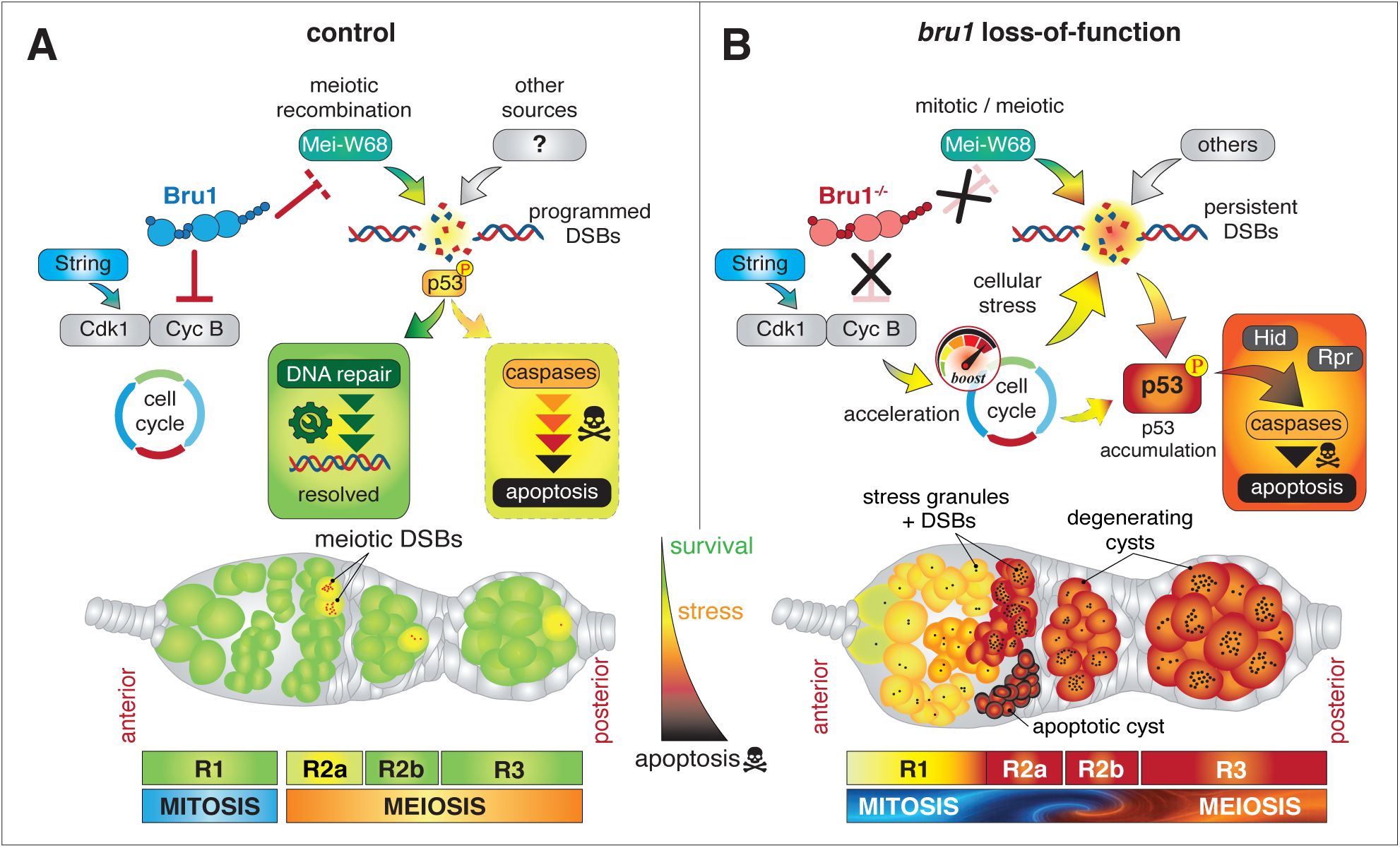
A model for Bru1 function in the germarium. (**A**) In control conditions, Bru1 regulates mitotic cyclin levels in region 1, and during the mitosis-to-meiosis transition in region 2. Germline proliferation progresses without noticeable DSBs. Upon meiotic DSBs induction, the limited breaks per cell are repaired and p53-dependent apoptosis is rarely activated. In our model, Bru1 acts as a break that permit the timely DNA repair contributing to the lack of DNA breaks. (**B**) In the absence of *bru1* function, Mei-W68 levels in germ cells are increased, generating non-meiotic DSBs in mitotically active region 1 cells and an excess of meiotic-DSBs in region 2. As a consequence, persistent DSBs are generated without being properly repaired. These unrepaired DSBs are accumulated and promote p53-mediated apoptosis, through the expression of *rpr* and *hid*. In addition, cyclins are not efficiently regulated and the mitotic cell cycle is accelerated, generating cellular stress(es), overwhelming the repair machinery and enhancing the DNA damage-mediated apoptosis of mutant cysts. Thus, the transition from mitosis to meiosis is not well stablished and oogenesis is blocked.

The absence of *bru1* function(s) in the germline results not only in altered cell cycles, but also in elevated *mei-W68 mRNA* and protein levels. In normal conditions, Mei-W68 generates DSBs in the pro-oocytes upon completion of the synaptonemal complex in late region 2a (Mehrotra & McKim, 2006). In contrast, *bru1* deficient cysts also exhibit Mei-W68-dependent DSBs in region 1. Since the latter occur prior to meiotic prophase I and since DSB repair in the female germline involves the homologue chromosome (Szostak *et al*, 1983; Haber, 2000), we speculate that DSBs generated in *bru1* mutant conditions cannot be as efficiently repaired in region 1 as they are in region 2 meiotic cells, in which homologues are paired. In addition, the increased amounts of Mei-W68 since early in oogenesis, and particularly in meiotic region 2, could imply a much higher frequency of DNA breaks, adding to the problem of break repair before oocyte specification. Thus, DSBs accumulate so that mutant region 1 and 2 cysts show precocious, excess breaks that eventually can cause cell death.

### *bru1* regulates DNA damage and stress granule formation

As a stress response, cells form reversible biomolecular condensates containing uncoated RNAs and RNA-binding proteins to protect mRNAs from damage and prevent the accumulation of incompletely translated nascent polypeptides (Reviewed in (Choi *et al*, 2025)). DNA damage can induce stress granule (SG) formation as an essential mechanism to sequester non-translating mRNAs and to regulate protein translation (Rhine *et al*, 2023). It is known that mitotic cyclin mRNAs (*CycA* and *CycB*) accumulate in P-bodies, where mRNAs are stored in a translationally repressed state and where, importantly, Bru1 is also present (Bayer *et al*, 2025; Milano *et al*, 2026). According to our model, the mRNAs of mitotic cyclins might be inefficiently protected in *bru1* loss-of-function conditions, hence preventing proper cyclin levels and justifying the unexpected mitotic cycles observed in the germline. Although accelerated cell cycle might prevent stress granules formation, as reported by other authors (Yahya *et al*, 2021), our observation of increased DNA damage and Caprin aggregates in *bru1* mutants suggest an alternative mechanism. Cell cycle alteration in *bru1* mutant region 1 might induce a high level of cellular stress such as oxidative stress, ER stress and/or DNA damage that may be reflected in a cumulative effect that peaks in region 2 and promotes stress granules formation. Since these phenotypes are rescued by depleting *string* quantities, the accumulation of stress granules observed under the mutant condition likely reflects an imbalanced cell cycle progression and the activation of the cellular stress response.

### *bru1* protects germ cells from meiotic DNA damage-induced apoptosis

The fact that programmed meiotic DSBs activate p53 without triggering apoptosis (Lu *et al*, 2010; Wylie *et al*, 2014) raises the important question of how germ cells are protected from the proapoptotic activity of p53. In somatic epithelial cells, DNA damage–induced proapoptotic activity of p53 is regulated by the G2/M-promoting factor Cdk1, such that in the absence of active Cdk1, p53 cannot induce cell death when the DNA repair mechanisms are operational (Ruiz-Losada *et al*, 2022; Qi & Calvi, 2016; Baonza *et al*, 2022). In this context, and since the downregulation of Cdk1 activity through Bru1-dependent degradation of mitotic cyclins is likely a key step in the mitosis-to-meiosis transition (Parisi *et al*, 2001; Sugimura & Lilly, 2006; Jones, 2004), we propose that *bru1* regulates the cell cycle to prevent active p53 acting in meiotic cells containing DNA damage. Thus, in the absence of *bru1*, Cdk1 activity is not properly silenced and cysts re-enter the mitotic cycle while activated p53 is dealing with meiotic DNA damage. The combination of both results in apoptosis, as reported for mouse gonads (Adhikari *et al*, 2016).

### Contribution of other Bru-like proteins to gametogenesis

While the involvement of a translational repressor such as Bru1 in the mitosis-to-meiosis transition may be a particularity of Drosophilids, post-transcriptional regulation seems to be a widespread strategy to ensure a swift transition. Entry into meiosis in mouse germ cells coincides with the pre-meiotic S phase, followed by a prolonged G2/meiotic prophase I, enabling meiosis-specific events such as chromosome synapsis and recombination to take place. Interestingly, while no specific translational repressor has been identified yet, some post-transcriptional regulators that are involved in the transition (e.g., MEIOC together with YTHDC2 and RBM46) act on mRNA stability to boost the expression of mitotic repressors thus enhancing an efficient activation of the meiotic programme (reviewed in (Ishiguro, 2023)).

In humans and mice, Bru-like proteins are a family of six RNA-binding proteins, CELF-1 to -6, which function in post-translational regulation including RNA stability and splicing (reviewed in (Vlasova-St. Louis *et al*, 2013)). Evidence from mice demonstrates that Bruno orthologues have important functions during gametogenesis or germ cell development. Thus, loss of CELF1 results in spermatogenesis and fertility defects in males and in reduced fertility in females, and CELF-3 mutant males display reduced sperm count and motility (Dev *et al*, 2007; Kress *et al*, 2006). In non-mammalian systems such as *Xenopus laevis*, one CELF orthologue is maternally expressed, suggesting a potential role in female gametogenesis (Wu *et al*, 2009). Altogether, these findings point to evolutionary conserved contributions of *bruno*-like genes in germline development, although defining their precise functions demands further work.

## Material & Methods

### *Drosophila* socks and maintenance

Flies were reared on standard wheat flour-agar medium at 25°C with relative humidity of approximately 50%. We used the following fly stocks:

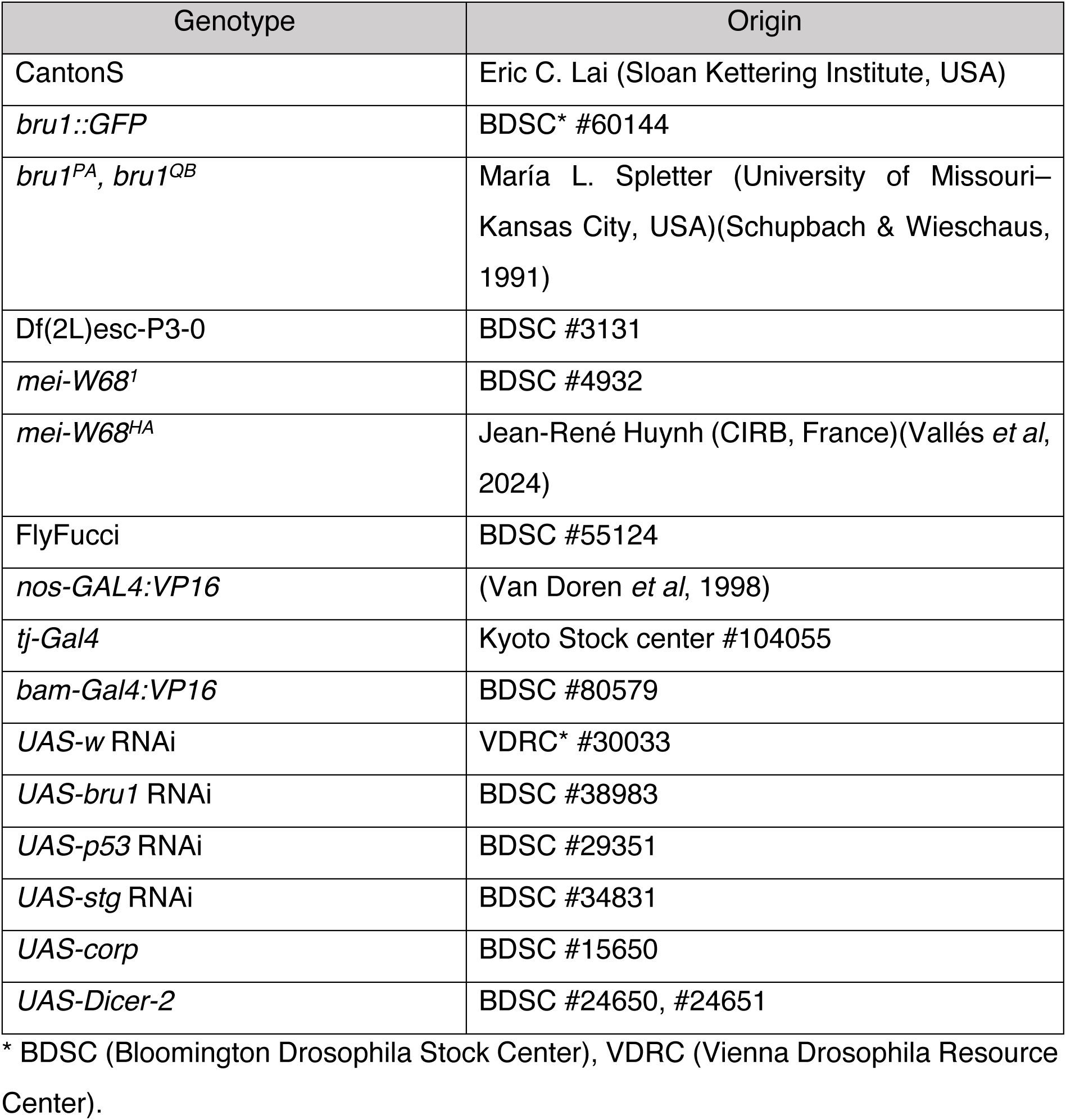

The CABD has authorization from the relevant authorities to use *Drosophila melanogaster* in laboratory research.

### Genotypes within figures

Figure 1:

B: CantonS
C: Bru1::GFP: y^1^ w^67c23^; Mi{PT-GFSTF.1}bru1^MI00135-GFSTF.1^

Figure 2:

control: CantonS
*bru1^PA/QB^*: *bru1^PA^ cn^-^ bw^-^/bru1^QB^ cn^-^ bw^-^*

Figure 3:

control: CantonS
*bru1^PA/QB^*: *bru1^PA^ cn^-^ bw^-^/bru1^QB^ cn^-^ bw^-^*

Figure 4:

control: CantonS
*mei-W68^1^*: *mei-W68^1^/mei-W68^1^*
*bru1^PA/QB^*: *bru1^PA^ cn^-^ bw^-^/bru1^QB^ cn^-^ bw^-^*
*bru1^PA/QB^ mei-W68^1^*: *bru1^PA^ cn^-^(?) mei-W68^1^/bru1^QB^ cn^-^(?) mei-W68^1^*
*mei-W68^HA/+^*: *mei-W68^HA^/+*
*bru1^PA/QB^ mei-W68^HA/+^*: *bru1^PA^ cn^-^(?) mei-W68^HA^/bru1^QB^ cn^-^ bw^-^*

Figure 5:

*nos>stgRNAi: UAS-Dicer-2/+; nos-Gal4/UAS-stg RNAi*
*nos>bru1^RNAi^, w^RNAi^: UAS-Dicer-2/UAS-bru1 RNAi;*
*nos-Gal4/UAS-w RNAi nos>bru1^RNAi^, stg^RNAi^: UAS-Dicer-2/UAS-bru1 RNAi; nos-Gal4/UAS-stg RNAi*

Figure 6:

*nos>wRNAi:* UAS-Dicer-2/+; nos-Gal4/UAS-w RNAi
*nos> bru1^RNAi^, w^RNAi^*: *UAS-Dicer-2/UAS-bru1 RNAi; nos-Gal4/UAS-w RNAi*
*nos>bru1^RNAi^, p53^RNAi^: UAS-Dicer-2/UAS-bru1 RNAi; nos-Gal4/UAS-p53 RNAi*
*nos>bru1^RNAi^, corp: UAS-Dicer-2/UAS-bru1 RNAi; nos-Gal4/UAS-corp*
*nos>bru1^RNAi^, hid^RNAi^: UAS-Dicer-2/UAS-bru1 RNAi; nos-Gal4/UAS-hid RNAi*
*nos>bru1^RNAi^, miRHG: UAS-Dicer-2/UAS-bru1 RNAi; nos-Gal4/UAS-miRHG*
control: CantonS
*bru1^PA^/bru1^QB^: bru1^PA^ cn^-^ bw^-^/bru1^QB^ cn^-^ bw^-^*

Figure S2:

Examples of egg chambers within *bru1^PA^ cn^-^ bw^-^/bru1^QB^ cn^-^ bw^-^* ovaries

Figure S3:

+/Df: +/Df(2L)esc-P3-0
*bru1^PA^/Df: bru1^PA^ cn^-^ bw^-^/*Df(2L)esc-P3-0
*bru1^QB^/Df: bru1^QB^ cn^-^ bw^-^/*Df(2L)esc-P3-0
control: CantonS
*bru1^PA/QB^*: *bru1^PA^ cn^-^ bw^-^/bru1^QB^ cn^-^ bw^-^bru1^-/-^*: *bru1^PA/QB^*, *bru^PA^/Df* or *bru^QB^/Df*

Figure S4:

control: CantonS
*bru1^PA/QB^: bru1^PA^ cn^-^ bw^-^/bru1^QB^ cn^-^ bw^-^*
*bru1^PA/QB^ mei-W68^1^: bru1^PA^ cn^-^(?) mei-W68^1^/bru1^QB^ cn^-^(?) mei-W68^1^*
*mei-W68^HA/+^*: *mei-W68^1^/+*
*bru1^PA/QB^ mei-W68^HA/+^*: *bru1^PA^ cn^-^(?) mei-W68^HA^/bru1^QB^ cn^-^ bw^-^*

Figure S5:

*nos>wRNAi:* UAS-Dicer-2/+; nos-Gal4/UAS-w RNAi
nos>bru1^RNAi^: UAS-Dicer-2/UAS-bru1 RNAi; nos-Gal4/+
tj>w^RNAi^: tj-Gal4/+; UAS-Dicer-2/UAS-w RNAi
tj>bru1^RNAi^: tj-Gal4/UAS-bru1 RNAi; UAS-Dicer-2/+
bam>w^RNAi^: bam-Gal4/+;; UAS-w RNAi
*bam>bru1^RNAi^: bam-Gal4; UAS-bru1 RNAi*

### *bru1* mutant alleles

EMS-induced mutations of *bru1*, including PA62 and QB72 and henceforth referred to as PA and QB, were first described as affecting spermatogenesis and oogenesis (Schupbach & Wieschaus, 1991; Webster *et al*, 1997). We established the presence of the PA and QB mutations in our fly stocks by genomic sequencing (Fig. S3A). The PA mutation causes an amino acid substitution in an RNA recognition motif, whereas the QB mutation results in the truncation of the protein removing the last RNA recognition motif (Fig. 1D) (Webster *et al*, 1997; Schupbach & Wieschaus, 1991; Parisi *et al*, 2001). We also confirmed that *bru1* was required for fertility, both in males and females (Fig. S3B). Mutant females showed severe defects in oogenesis affecting early egg chambers and completely abrogating egg production. Depending on the strength of the allelic combination, oogenesis could proceed until mid-oogenesis (stages (S) 8-9) before degenerating (*bru1^PA^*/*Df(2L)esc-P3-0*), or it could be disrupted from its earliest stages, yielding almost no egg chambers (*bru1^QB^*/*Df(2L)esc-P3-0*). The *bru1^PA^*^/*QB*^ trans-heterozygous combination also failed to produce eggs but exhibited milder ovariole degeneration compared to *bru1^PA^*/*Df(2L)esc-P3-0* hemizygous mutants (Fig. S3C-F).

### DNA extraction and amplification

1-3 flies were ground twice with a pestle for 2 min adding 200 *µ*L lysis buffer (100 mM TrisHCl pH 7.5, 100 mM EDTA, 100 mM NaCl, 0.5% SDS). The lysate was incubated at 65°C for 30 min. The DNA was obtained following a standard LiCl/KAc-isopropanol protocol. The DNA pellet was dried and resuspended in 50 *µ*L TE buffer (10 mM TrisHCl pH 8.0, 1 M EDTA). DNA was amplified using a standard PCR protocol. ∼100 ng of DNA per reaction were amplified using BioTaq^TM^ DNA polymerase (BIOTLINE, BIO-21040). We used the following primers:

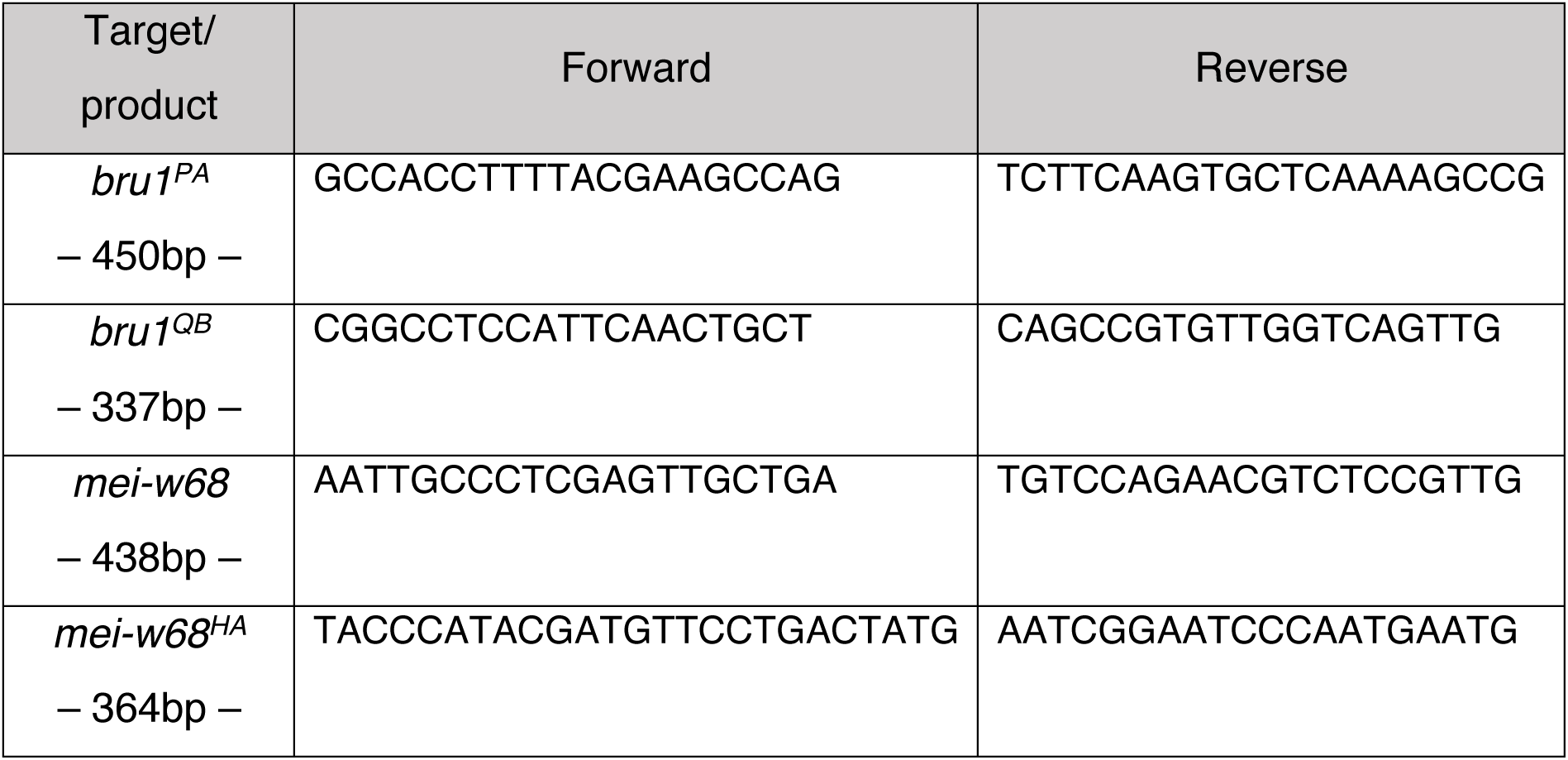

### Generation and detection of recombinants

To generate *bru1^PA^* (or *bru1^QB^*) and *mei-w68^1^* recombinants, we used initially the crushed thorax phenotype as a marker for the presence of the *bru1* mutations, as previously described (Spletter *et al*, 2015). To confirm molecularly the presence of *bru1* mutations, DNA from hemizygous flies (recombinant chromosome/ Df (2L) esc-P3-0) was amplified and sequenced to detect the *bru1^PA^*or *bru1^QB^* mutations. Both mutations generated nucleotide changes (*bru1^PA^*: A>C; …TGCTTTGC**<u>A</u>**TAATGTTA…; *bru1^QB^*: C>T; …TGGGAGCC**<u>C</u>**AGCAGAGC…). To check for the *mei-w68^1^* mutation, genomic DNA from putative homozygous flies was amplified by PCR. Since the *mei-w68^1^* mutation is caused by a deletion in the gene and because of the primers used, the mutation was revealed by the absence of PCR product.

To generate the *bru1^PA^* and *mei-w68^HA^*recombinant, we followed a similar strategy. *mei-w68^HA^* contains an 3′ HA-tag (3xHA-GTTAAA-6XHis) that can be detected by PCR or with an anti-HA antibody (Roche, #11867423001). Of note, the HA insertion in *mei-W68* compromises its functionality and causes subfertility in homozygosis (Vallés *et al*, 2024). In our experiments, we worked in heterozygous conditions.

### Fertility assays

Fertility was measured using a binary assay. 10 individual crosses were set with single males and females for each genotype. Couples were placed in tubes with fresh fly food and transferred to new food every 3 days. All tubes were monitored for 14 days. We scored the presence of eggs, larvae, pupa and/or offspring.

### Immunofluorescence staining and image acquisition

Ovaries from 5-10 yeast-fed females were dissected on PBS 1x + 0.1% Triton-X100. While dissecting, the collection tube was kept on ice. Ovaries were fixed on a rotator for 20 min in 4% formaldehyde + 0.1% Tween-20. After washing the fixative, ovaries were permeabilized for 1 hour using 1% Triton-X100 and then blocked for 1 extra hour using PBT_10_+NP-40 (PBT + 10% BSA + 0.1% NP-40). Primary antibodies were incubated in PBT_1_ (PBT + 1% BSA) overnight at 4°C. After washing the primary antibodies, secondary antibodies were incubated 1:100 in PBT_0.1_ (PBT + 0.1% BSA) for 4 hours. After incubation and washing, ovaries were incubated, when required, with 1:200 Rhodamine: Phalloidin and 1:500 Hoechst for 30 min. Ovaries were mounted in antifade mounting media (VectaShield®).

Images were acquired with an inverted confocal microscope (Leica Stellaris 5, model DMI8) using a 63x/1.4 NA oil-immersion, planApo-corrected, objective. Depending on the fluorophores used, samples were excited in two tracks by combining laser lines as follows: 405+561 and 488+638. Typically, images were acquired at 0.5 *µ*m intervals and with a digital zoom between 0.75x and 2x. Images were analyzed using ImageJ (Fiji).

List of primary antibodies and dyes used:

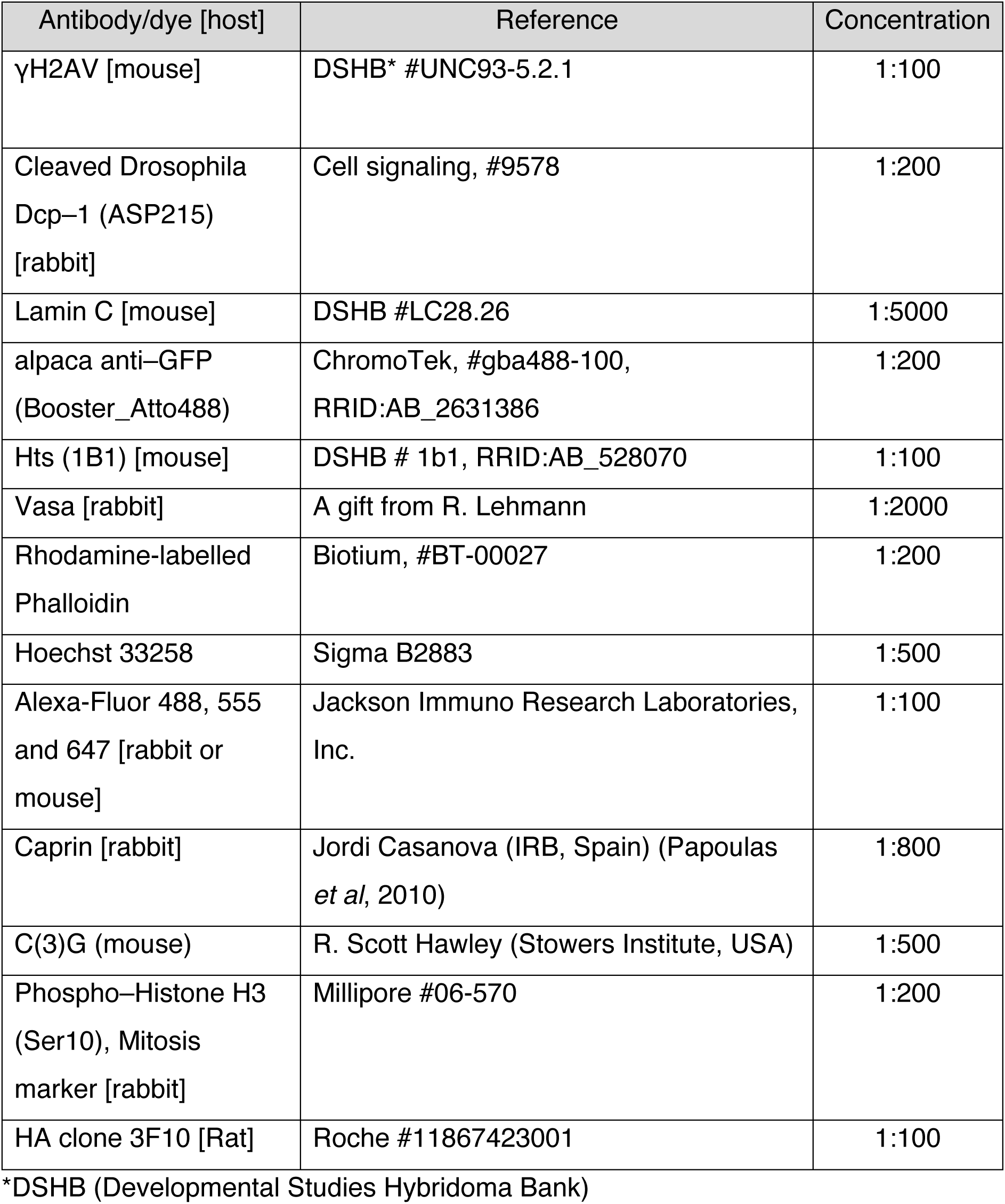

### Quantification of germ cell phenotypes

The different germ cell types in the germarium (GSCs, CBs, 2-16 cell cysts) were scored based on the visualization of different markers such as the spectrosome/fusome (anti-Hts), nucleus (anti-Lamin C or Hoechst) and cytoplasm (anti-Vasa) and their localization within the niche. GSCs were identified by their spectrosome figures (Villa-Fombuena *et al*, 2021) and by their attachment to Cap cells (CpCs). Single germ cells detached from CpCs and containing a spectrosome were classified as CBs. 2-16 cell cysts contained fusomes of distinct morphologies. Nuclear sizes were also useful to determine whether cells belonged to the same cyst. Region 3/S1 cysts are spherical and remain at the posterior of the germarium. S2-5 were scored based on their whole size and on particular characteristics of each stage (see more details in Figure 1 from (Jia *et al*, 2016)). Mutant cysts and egg chambers could show abnormal numbers of germ cells. We confirmed the presence of 32cc and 64cc by measuring nuclear size. Encapsulation defects were detected in egg chambers containing fewer than 16 cystocytes, and these underloaded egg chambers were frequently adjacent to others containing the rest of the full 16-cell complement. Fused egg chambers were discarded from our quantifications.

### Quantification of S-phase and mitotic cells

To score S-phase cells, 5-ethynyl-2^0^-deoxyuridine (EdU) labeling was used with the ‘‘Alexa Fluor 647 dye, Click-iT EdU Cell Proliferation Kit for imaging’’ (Thermo Fisher Scientific, Cat# C10640). Dissected samples were incubated for 30 min in Schneider’s medium (Biowest, #L0207-500) supplemented with 10mM EdU prior to fixation. Samples were then incubated during 30 min for the click chemical reaction following the manufacturer’s instructions. To quantify mitotic cells, we used an antibody against the phosphorylated histone H3 at serine 10 (Millipore, #06-570), a marker of chromatin condensation during mitosis. Data were plotted as the number of PH3^+^ cells divided by the total number of cysts in the germarium. For either of these analyses, more than 20 germaria were studied per each condition and replicate.

### Quantification of Flyfucci signal

The Flyfucci strategy (Zielke *et al*, 2014) uses GFP- and RFP-tagged degrons from E2F1 and Cyclin B proteins, respectively, to label G1 (only GFP::E2F1 is expressed), S (only RFP::CycB) and G2/M (both GFP::E2F1 and RFP::CycB). To quantify the Flyfucci signal, we established the nuclear area as determined by the Hoechst marker to measure the RFP and GFP nuclear signals. We stablished a detection threshold of 13 arbitrary units (a.u.) for the RFP or GFP values so that signals <13 were considered as negative. Using RFP and GFP intensity values, we calculated RFP/GFP ratios to categorize cells as red (ratio >1.9), green (ratio <0.7) and yellow (0.7 <ratio <1.9).

### Quantification of γH2AV, Caprin aggregates, apoptosis and HA levels

Ovaries stained with the anti-γH2AV antibody (Lake *et al*, 2013), were quantified as follows. Nuclear area was delimited using the Hoechst staining and γH2AV intensity was then obtained and plotted. We scored 100-500 nuclei per germarial region or stage from ≈20 germaria (each genotype: 500-1500 single values). anti-Vasa staining was used to confirm germline origin. Medians (line) and means (cross) are shown in violin plots.

To quantify the number of Caprin aggregates, we used Fiji (ImageJ) plugins to acquire z-projections, segment the signal and count particles. The number of Caprin aggregates was scored in early/late region 1 and regions 2a, 2b and 3. Results were expressed as the number of aggregates per area unit (*µ*m²).

Ovaries immunostained with anti-Cleaved-Dcp1 (Cell signaling, #9578) were quantified in a binary manner (presence or absence of Dcp1 activation), using a threshold established by comparing intensity values of Dcp1^+^ vs Dcp1^-^ cysts. 40<x<130 germaria were analyzed.

To quantify the HA signal, multiple areas in regions 1, 2a, 2b and 3 were selected to obtain mean HA values. 10<x<20 germaria were analyzed.

### HCR RNA-FISH

Hybridization Chain Reaction (HCR) RNA fluorescent in situ hybridization was performed on *Drosophila* ovaries using HCR™ RNA-FISH bundles (Molecular Instruments) following the manufacturer’s protocol with minor modifications. Probe sets were used to target *rpr* (amplifiers B1-546), *hid* mRNA (amplifiers B2-488) and *mei-W68* mRNA (amplifiers B1-488). Ovaries were dissected in Schneider’s medium with 10% FBS and fixed in 4% paraformaldehyde containing 0.1% sodium deoxycholate and 0.3% Triton X-100 for 1 hour at RT. Samples were washed once in PBS for 10 min, permeabilized with PBTw (PBS + 0.1% Tween-20) for 30 min and incubated with pre-warmed hybridization buffer (300 *µ*L/sample) at 37 °C for 30 min. 1 pmol of each probe in 100 *µ*L hybridization buffer (1 *µ*L/sample of rpr-B1 and hid-B2) were added directly to the samples and incubated overnight at 37 °C with gentle agitation. After hybridization, samples were washed 4×15 min with pre-warmed probe wash buffer (37 °C) and 2×5 min with sodium chloride-sodium citrate (SSC) 5× supplemented with 0.1% Tween (SSCT) at RT. Next, samples were incubated at RT for 30 min in 300 *µ*L amplification buffer. 2 *µ*L from each hairpin/sample (3 *µ*M stocks: B1-H1-488, B1-H2-488, B2-H1-546 and B2-H2-546) in 100 *µ*L amplification buffer were snap-heated at 95°C for 90 sec, preserved protected from light at RT for 30 min, added the samples and incubated O/N at 25 °C in the dark with gentle shaking. Next day, ovaries were washed sequentially with 5× SSCT (2×5 min, 2×30 min), incubated with Hoechst (1:100) for 30 min, washed 3×10 min in PBS, and mounted in VectaShield for imaging. To quantify the number of mRNA foci, we used Fiji (ImageJ) plugins to acquire z-projections, segment the signal and count particles. The number of *rpr* or *hid* mRNA foci was scored in germarial regions 1, 2 and 3. Results were expressed as the number of mRNA foci per area unit (*µ*m²).

### Plotting data and statistical analyses

Experiments were performed with at least three biological replicates. Samples were collected from at least 5 different individuals grown in equivalent conditions. For each quantification (except in specified cases), more than 20 germaria were analyzed per condition and per replicate. For violin graphs, data plotted for each genotype represents 500-1500 single values. These graphs show the median (line) and mean (X). Scatter plots and bar graphs indicate mean±SEM. Sample sizes shown in each panel correspond to the number of germaria or egg chambers analyzed. Statistically significant differences between control and experimental conditions were calculated with unpaired Student’s, nonparametric Mann-Whitney, or Chi-square t-tests, as indicated (*=p<0.05; **=p<0.01; ***=p<0.001; ****=p<0.0001). Only significant differences are indicated in the graphs.

## Acknowledgements

This work was supported by the Spanish Agencia Estatal de Investigación (MCIU/AEI, http://www.ciencia.gob.es/; grant numbers PID2021-125480NB-I00 and PID2024-155234NB-100 to AG-R; and PID2021-127114NB-I00 and PID2024-160216NB-I00 to CE), and by the Talent Attraction program from the Junta de Andalucía (grant number P21_00450 to JG-M). We thank Jordi Casanova for his support. We are grateful to María L. Spletter for kindly providing the *bru1^PA^* and *bru1^QB^*fly stocks. We are also grateful to Lola Martín-Bermudo and Salvador Herrera and their teams for constructive discussions.

## Author contributions

Conceptualization, CE & AG-R; Methodology, JG-M; Formal analysis, JG-M & AG-R; Investigation, JG-M; Resources, AG-R; Writing – Original draft, JG-M, CE & AG-R, Writing – Review & Editing, JG-M, CE & AG-R &; Project administration, JG-M; Visualization, JG-M; Supervision, CE & AG-R; Funding Acquisition, AG-R.

## Supplemental information titles and legends

Document S1. Figures S1-S5.

## SUPPLEMENTARY FIGURE LEGENDS

**Figure S1.**
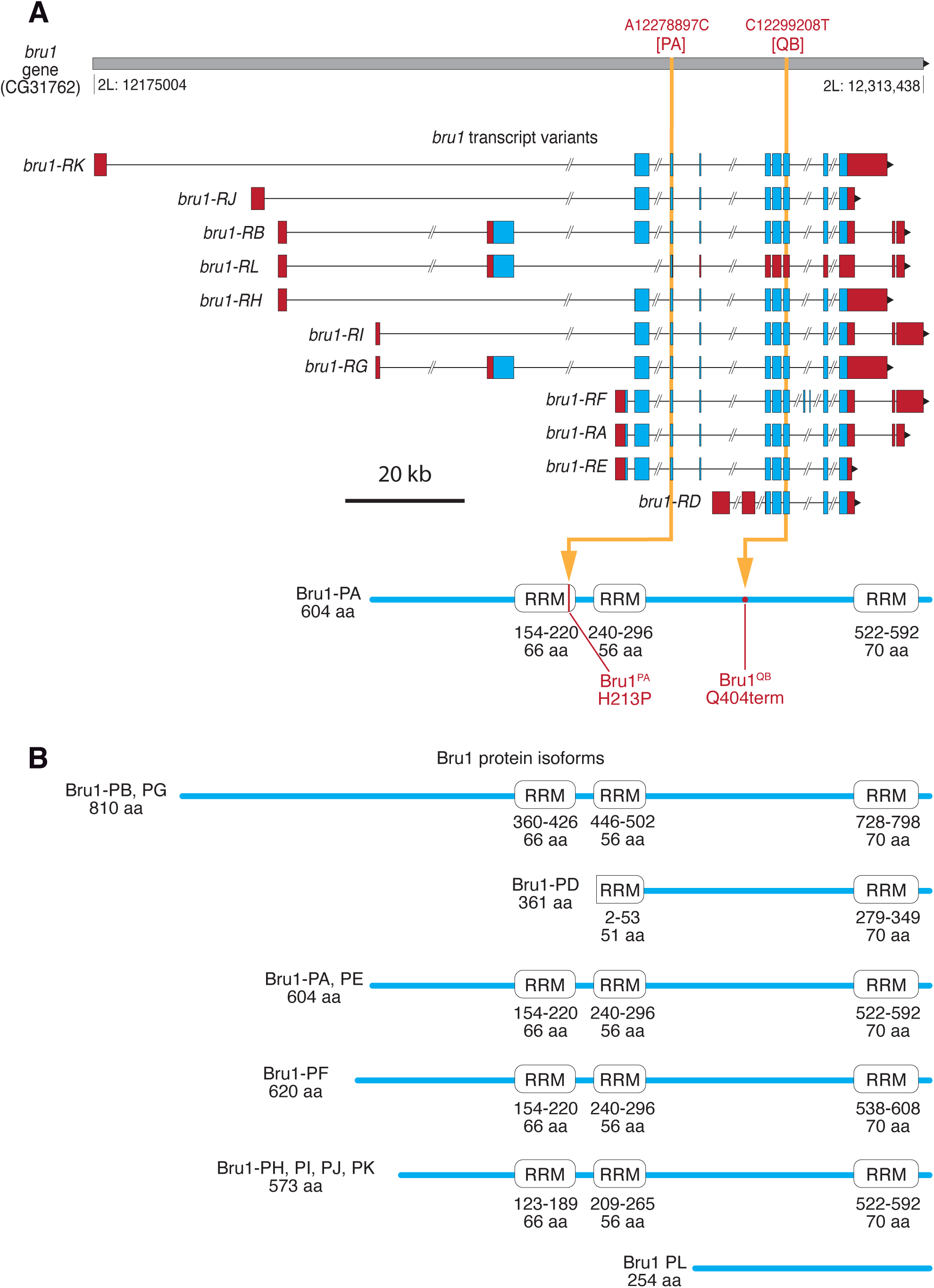
Transcripts and protein isoforms of *bru1*. (**A**) Scheme of the *bru1* locus highlighting the predicted transcript variants. Untranslated regions (UTRs) are shown in red, and open reading frames (ORFs) are shown in blue. Among the *bru1* transcripts variants, a representative protein isoform (isoform A) is shown. Bru1-PA contains three RNA recognition motifs (RRMs). Yellow arrows indicate the location of the PA and QB mutations in the transcripts and corresponding protein. The PA mutation causes an amino acid substitution (H213P) that affects the first RRM, whereas the QB mutation results in a truncated protein (Q404term), preventing translation of the third RRM. Genomic coordinates according to FlyBase are indicated (release FB2025_04). (**B**) Scheme of each of the predicted Bru1 protein isoforms.

**Figure S2.**
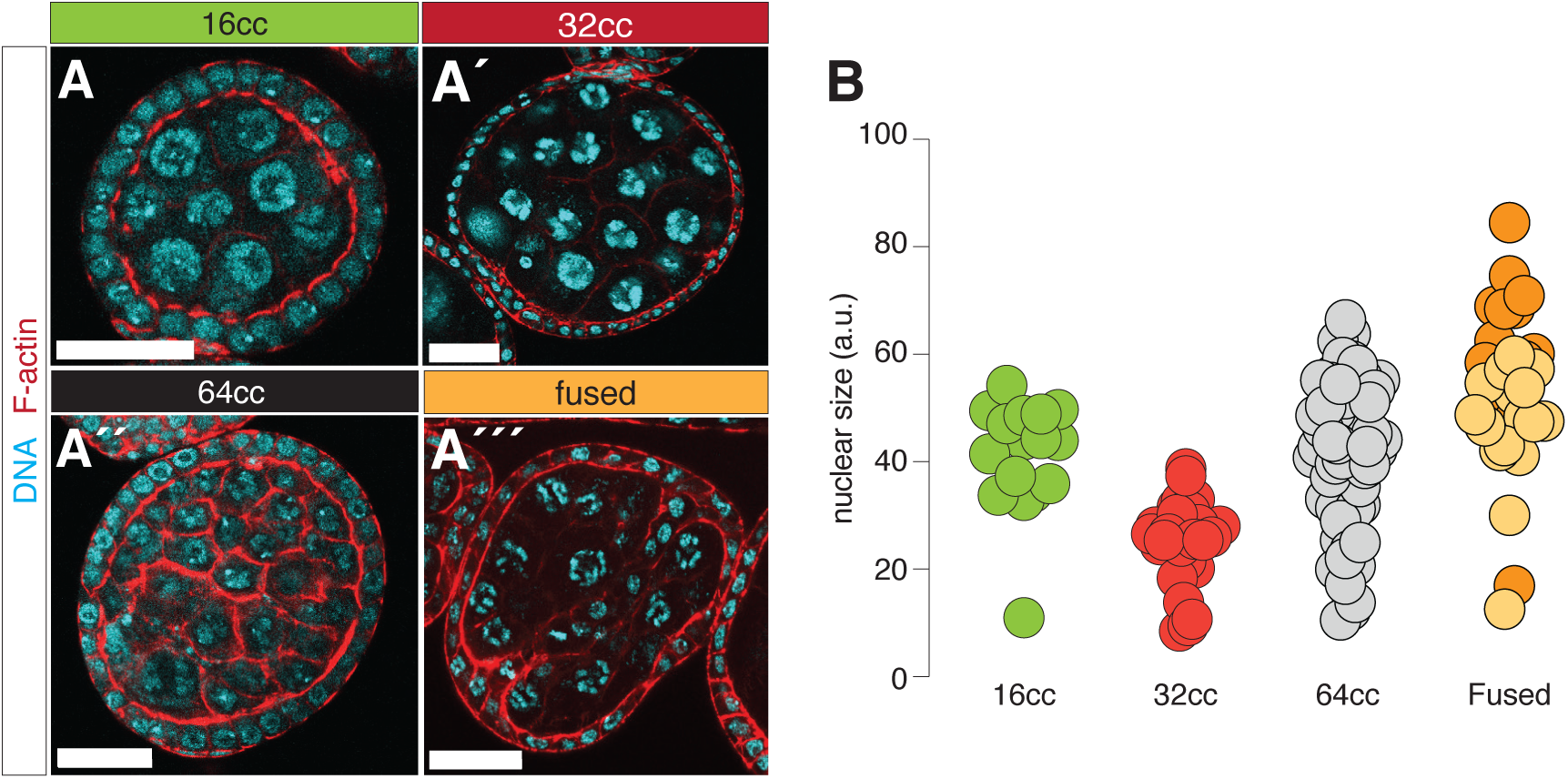
Examples of cyst with extra sets of germline cells found in *bru1* mutants. (**A**) Egg chambers stained to visualize DNA (Hoechst, cyan) and F-actin (Rhodamine: Phalloidin, red). Examples of control, 32cc, ≈64cc and fused egg chambers are shown. (**B**) The germline nuclear size can be used to distinguish fused egg chambers from those with extra divisions, as the former contain clusters of larger and smaller sizes whereas the latter are more homogeneous.

**Figure S3.**
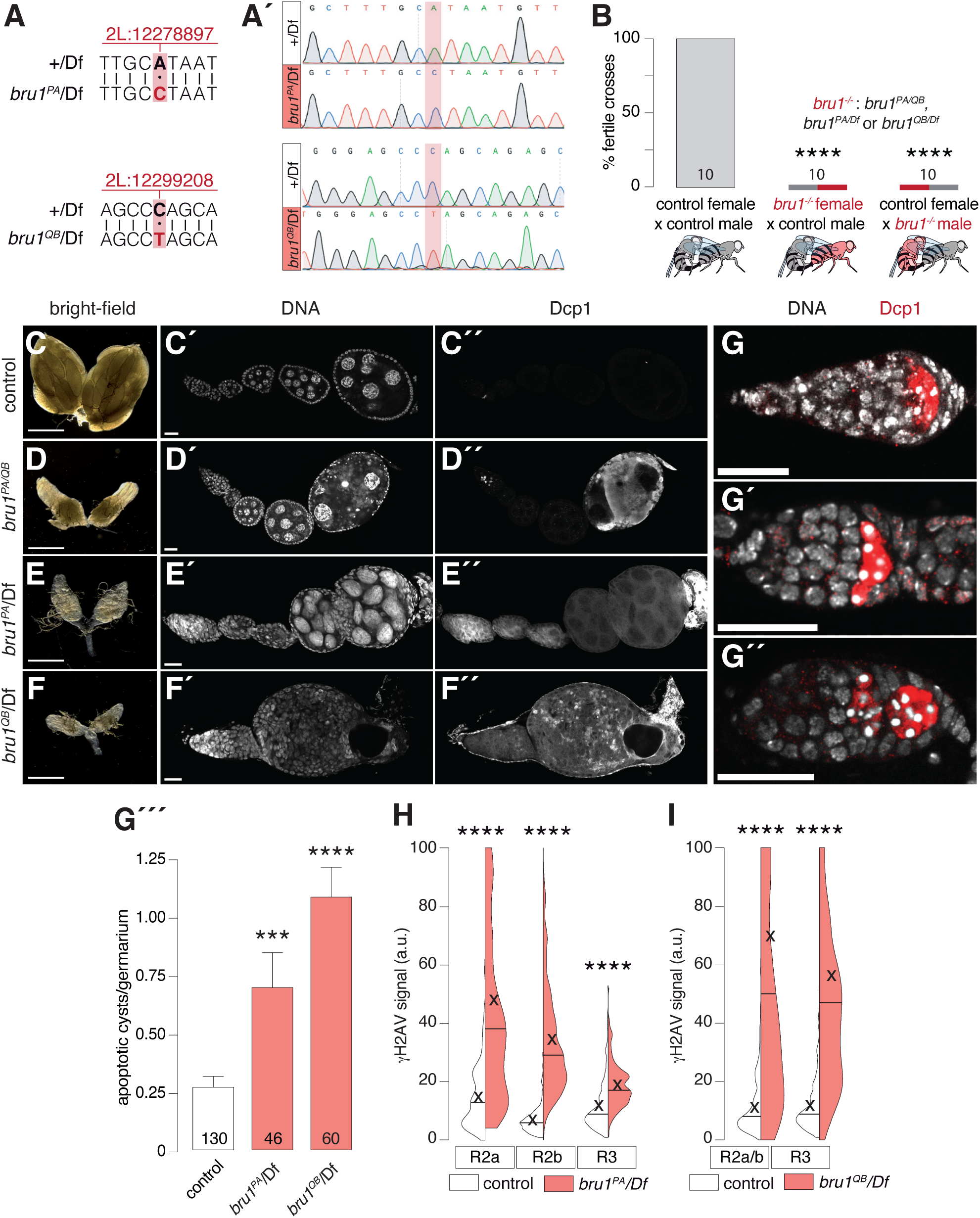
Hemizygotes phenocopy *bru1* loss of function phenotype. (**A**) Sequence confirmation of the presence of the PA and QB mutations. (**B**) Percentage of fertile crosses of control and *bru1^-/-^* mutant flies. (**C-F**) Dissecting scope images of ovaries. Ovarioles stained to visualize apoptotic cysts (anti-Dcp1) and DNA (Hoechst). (F) Examples of *bru1* mutant germaria showing apoptotic cysts (anti-Dcp1, red) presenting pycnotic nuclei (Hoechst, white). Quantification of the number of apoptotic cysts per germarium is shown in (G′′′). (**H**, **I**) Quantification of nuclear γH2AV signal in regions 2a, 2b and 3 of controls and *bru^PA^/Df* or *bru^QB^/Df*. Statistical significance was calculated with Chi-square (B), unpaired Student’s (G) and Mann-Whitney unpaired nonparametric t-tests (H, I). Scale bars: 500 μm for ovaries and 25 μm for ovarioles.

**Figure S4.**
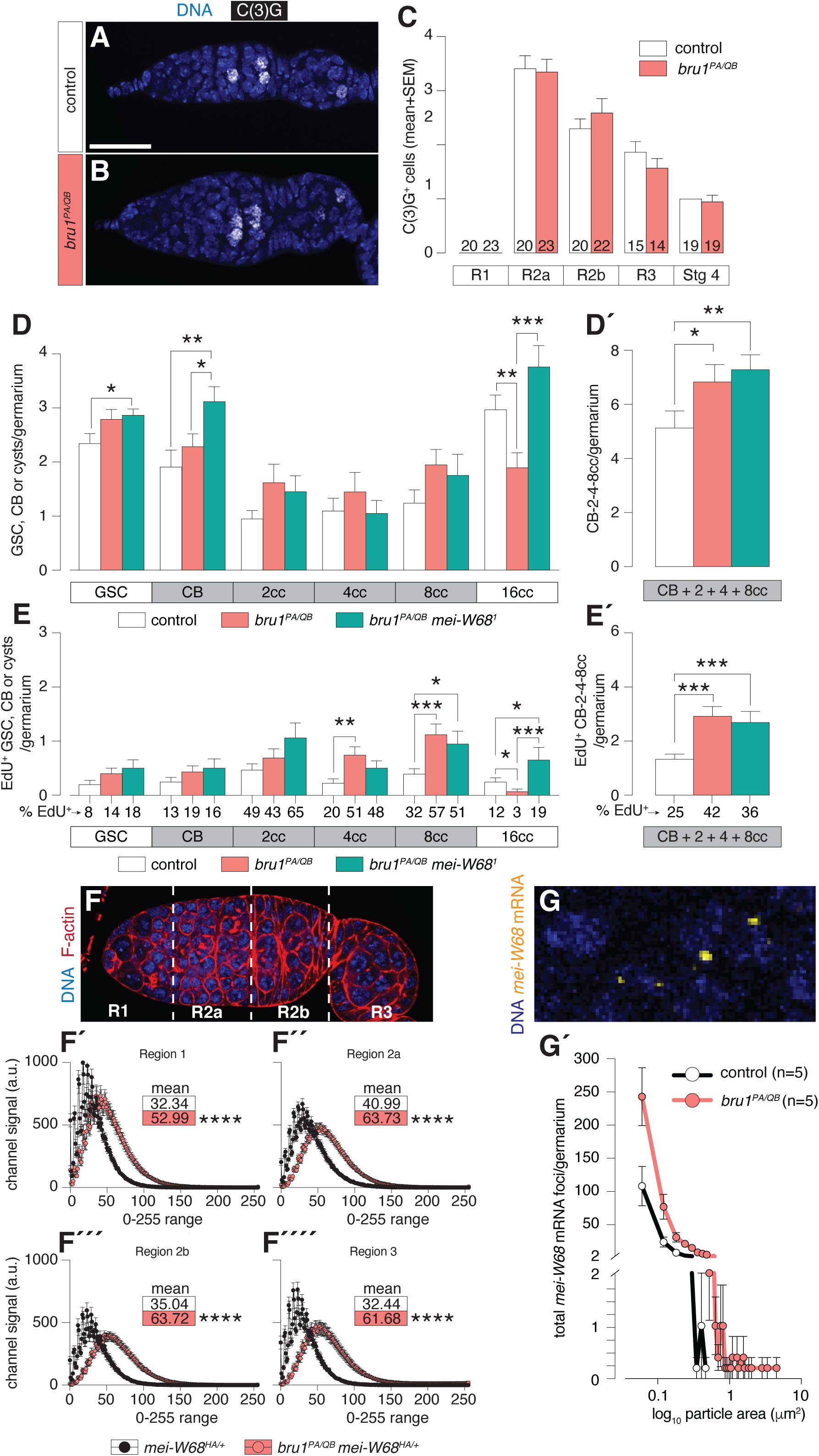
Supporting data for the cell cycle phenotypes from. **Figure 4**. (**A**, **B**) Germaria stained to visualize synaptonemal complex formation (C(3)G, white), and counterstained with DNA (Hoechst, blue). (**C**) Quantification of number of cells containing C(3)G^+^ figures per regions/stage. (**D**) Quantification of the total number of GSCs, CBs and 2-16 cell cysts. The total number of CBs + 2-8 cell cysts is shown in (D′). (**E**) Quantification of EdU^+^ GSCs, CBs and 2-16 cell cysts. The total number of EdU^+^ CBs + 2-8 cell cysts is shown in (É). (**F**) Germarium stained to visualize F-actin (Rhodamine: Phalloidin, red) and DNA (Hoechst, blue) delimiting regions 1, 2a, 2b and 3. HA signal levels were quantified per region and values plotted through 0-255 range scale. (**G**) Detail of germarium stained to visualize *mei-W68* mRNA (yellow) and DNA (Hoechst, blue). The number of foci/germarium *versus* particle size (log_10_) is plotted in (G′). Statistical significance was calculated with an unpaired Student’s t-test (A, B, C, F). Scale bars: 25 μm.

**Figure S5.**
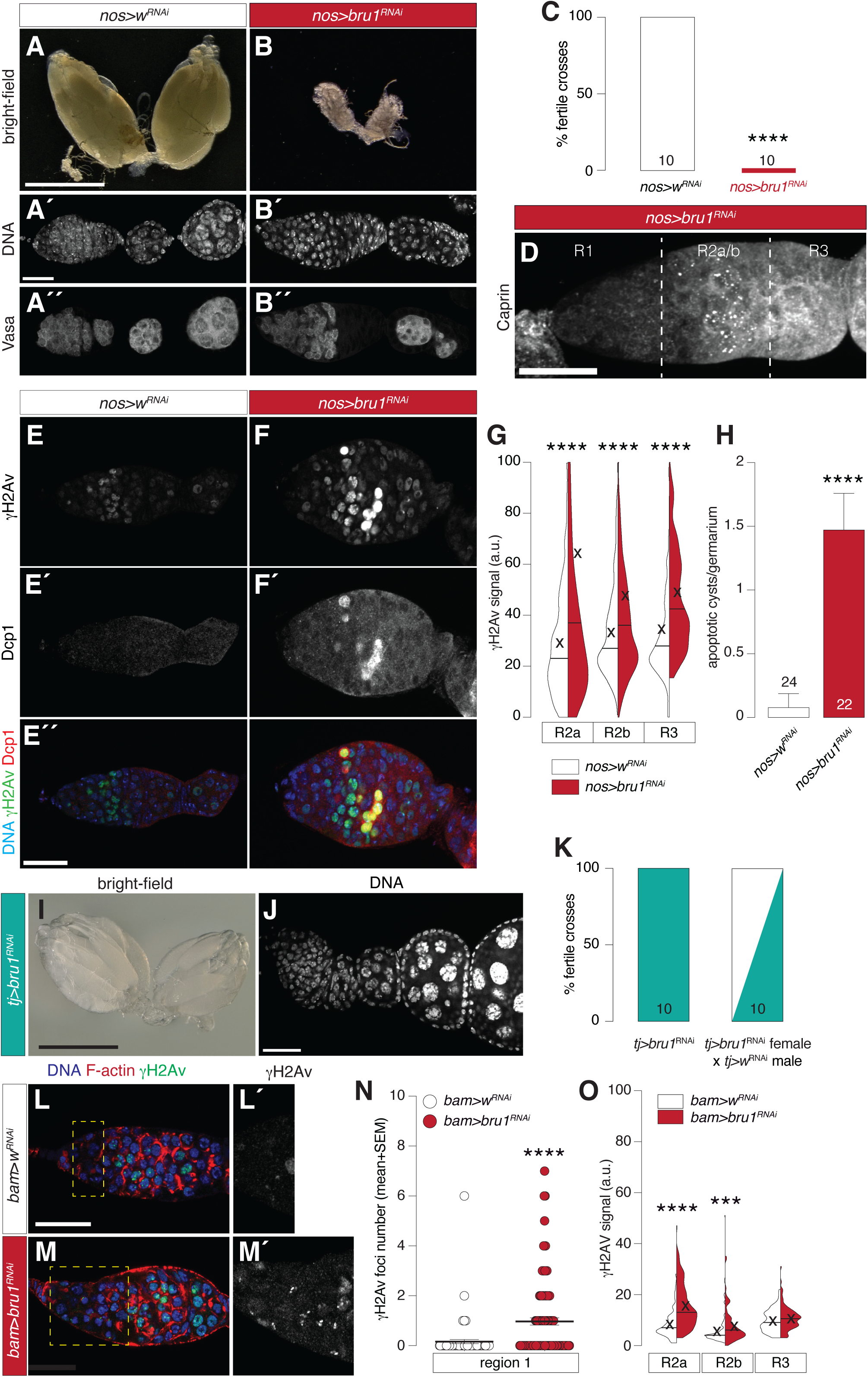
Downregulation of *bru1* in the germline mimics the *bru1* mutant phenotypes. (**A**, **B**) Dissecting scope images of *nos>w* RNAi and *nos>bru1* RNAi ovaries. Germaria stained to visualize germline cells (Vasa) and DNA (Hoechst). (**C**) Percentage of fertile crosses. (**D**) Germarium stained to visualize stress granules (anti-Caprin 1). (**E**, **F**) Germaria stained to visualize DSBs (anti-γH2AV, green), apoptotic cysts (anti-Dcp1, red) and DNA (Hoechst dye, blue). (**G**) Quantification of nuclear γH2AV signal in regions 2a, 2b and 3. (**H**) Quantification of the number of apoptotic cysts per germarium. (**I**, **J**) Downregulation of *bru1* in somatic cells using *traffic jam* (*tj*)-Gal4 as a driver. Images of an ovary and an ovariole stained with the DNA dye Hoechst (white) to visualize the structural integrity. (**K**) Percentage of fertile crosses using *tj>bru1* RNAi females and either *tj>bru1* RNAi and *tj>w* RNAi males. (**L, M**) Downregulation of *bru1* in region 1 (CBs and early cysts) using *bag-of-marbles* (*bam*)-Gal4 as a driver. Germaria stained to visualize DSBs (anti-γH2AV, green), F-actin (Rhodamine: Phalloidin, red) and DNA (Hoechst, blue). Insets show γH2AV detail in region 1. (**N**) Number of γH2AV foci in region 1 germline cells of 3 *bam*-driven *bru1* depleted mutant germaria. (**O**) Quantification of nuclear γH2AV signal in regions 2a, 2b and 3 upon *bam*-driven *bru1* depletion. Data plotted were collected from 7 germaria (each genotype: 600-700 single values). Statistical significance was calculated with Chi-square (C, K), Mann-Whitney unpaired nonparametric (G, N, O) and unpaired Student’s t-tests (H). Scale bars: 500 μm for ovaries and 25 μm for ovarioles.

